# Insights into Genome Recoding from the Mechanism of a Classic +1-Frameshifting tRNA

**DOI:** 10.1101/2020.12.31.424971

**Authors:** Howard Gamper, Haixing Li, Isao Masuda, D. Miklos Robkis, Thomas Christian, Adam B. Conn, Gregor Blaha, E. James Petersson, Ruben L. Gonzalez, Ya-Ming Hou

**Affiliations:** Department of Biochemistry and Molecular Biology, Thomas Jefferson University, Philadelphia, PA 19107, USA; Department of Chemistry, Columbia University, New York, NY 10027, USA; Department of Chemistry, University of Pennsylvania, Philadelphia, PA 19104, USA; Department of Biochemistry, University of California, Riverside, CA 92521, USA

**Author notes:** These authors contributed equally to this work (Howard Gamper, Haixing Li). Corresponding authors (T); (F); (T); (F). Lead contact: Ya-Ming Hou.

**Keywords:** *SufB2* frameshift suppressor tRNA, +1 ribosomal frameshifting, quadruplet codon, genome expansion, m^1^G37 methylation

## Abstract

While genome recoding using quadruplet codons to incorporate non-proteinogenic amino acids is attractive for biotechnology and bioengineering purposes, the mechanism through which such codons are translated is poorly understood. Here we investigate translation of quadruplet codons by a +1-frameshifting tRNA, *SufB2*, that contains an extra nucleotide in its anticodon loop. Natural post-transcriptional modification of *SufB2* in cells prevents it from frameshifting using a quadruplet-pairing mechanism such that it preferentially employs a triplet-slippage mechanism. We show that *SufB2* uses triplet anticodon-codon pairing in the 0-frame to initially decode the quadruplet codon, but subsequently shifts to the +1-frame during tRNA-mRNA translocation. *SufB2* frameshifting involves perturbation of an essential ribosome conformational change that facilitates tRNA-mRNA movements at a late stage of the translocation reaction. Our results provide a molecular mechanism for *SufB2*-induced +1 frameshifting and suggest that engineering of a specific ribosome conformational change can improve the efficiency of genome recoding.

## INTRODUCTION

The ability to recode the genome and expand the chemical repertoire of proteins to include non-proteinogenic amino acids promises novel tools for probing protein structure and function. While most recoding employs stop codons as sites for incorporating non-proteinogenic amino acids, only two stop codons can be simultaneously recoded due to the cellular need to reserve the third stop codon for termination of protein synthesis. The use of quadruplet codons as additional sites for incorporating non-proteinogenic amino acids has thus emerged as an attractive alternative^1,2^. Recoding at a quadruplet codon requires a +1-frameshifting tRNA that is aminoacylated with the non-proteinogenic amino acid of interest. The primary challenge faced by this technology has been the low efficiency with which the full-length protein carrying the non-proteinogenic amino acid can be synthesized. One reason for this is the poor recoding efficiency of the +1-frameshifting aminoacyl (aa)-tRNA, and the second is the failure of the +1-frameshifting aa-tRNA to compete with canonical aa-tRNAs that read the first three nucleotides of the quadruplet codon at the ribosomal aa-tRNA binding (A) site during the aa-tRNA selection step of the translation elongation cycle. While directed evolution by synthetic biologists has yielded +1-frameshifting tRNAs, efficient recoding requires cell lines that have been engineered to deplete potential competitor tRNAs^3–8^. These problems emphasize the need to better understand the mechanism through which quadruplet codons are translated by +1-frameshifting tRNAs.

In bacteria, +1-frameshifting tRNAs that suppress single-nucleotide insertion mutations that shift the translational reading frame to the +1-frame have been isolated from genetic studies^9,10^. These +1-frameshifting tRNAs typically contain an extra nucleotide in the anticodon loop – a property that has led to the proposal of two competing models for their mechanism of action. In the quadruplet-pairing model, the inserted nucleotide joins the triplet anticodon in pairing with the quadruplet codon in the A site and this quadruplet anticodon-codon pair is translocated to the ribosomal peptidyl-tRNA binding (P) site^11^. In the triplet-slippage model, the expanded anticodon loop forms an in-frame (0-frame) triplet anticodon-codon pair in the A site and subsequently shifts to the +1-frame at some point later in the elongation cycle^12,13^, possibly during translocation of the +1-frameshifting tRNA from the A to P sites^14^ or within the P site^15^. The triplet-slippage model is supported by structural studies of ribosomal complexes in which the expanded anticodon-stem-loops (ASLs) of +1-frameshifting tRNAs have been found to use triplet anticodon-codon pairing in the 0-frame at the A site^16–18^ and in the +1-frame at the P site^19^. Nonetheless, these structures do not eliminate the possibility that two competing triplet pairing schemes (0-frame and +1-frame) can co-exist when a quadruplet codon motif occupies the A site^15^, that some amount of +1 frameshifting can occur via the quadruplet-pairing model, and that the quadruplet-pairing model may even dominate for particular +1-frameshifting tRNAs, codon sequences, and/or reaction conditions^10^. We also do not know how each model determines the efficiency of +1 frameshifting or whether any competition between the two models is driven by the kinetics of frameshifting or the thermodynamics of base pairing. In addition, virtually all natural tRNAs contain a purine at nucleotide position 37 on the 3’-side of the anticodon (http://trna.bioinf.uni-leipzig.de/), which is invariably post-transcriptionally modified and is important for maintaining the translational reading frame in the P site^15^. While most +1-frameshifting tRNAs sequenced to date also contain a purine nucleotide at position 37 ^8^, we do not know whether it is post-transcriptionally modified or how the modification affects +1 frameshifting. Perhaps most importantly, while the structural studies described above provide snapshots of the initial and final states of +1 frameshifting, they do not reveal where, when, or how the shift occurs, thereby precluding an understanding of the structural basis and mechanism of +1 frameshifting. These open questions have limited our ability to increase the efficiency of genome recoding at quadruplet codons.

To address these questions, we have investigated the mechanism of +1 frameshifting by *SufB2* (Figure 1a), a +1-frameshifting tRNA that was isolated from *Salmonella typhimurium* as a suppressor of a single C insertion into a proline (Pro) CCC codon^20^. The observed high +1-frameshifting efficiency of *SufB2* at the CCC-C motif, nearly 80-fold above background^20^, demonstrates its ability to successfully compete with the naturally occurring *ProL* and *ProM* isoacceptor tRNAs that read the CCC codon. Using the ensemble ‘codon-walk’ methodology^21^ and single-molecule fluorescence resonance energy transfer (smFRET), we have compared the +1 frameshifting activity of *SufB2* relative to its closest counterpart, *ProL*, at a CCC-C motif, and determined the position and timing of the shift. Our results show that *SufB2* is naturally *N*^1^-methylated at G37 in cells, generating an m^1^G37 that blocks quadruplet pairing and forces *SufB2* to use 0-frame triplet anticodon-codon pairing to decode the quadruplet codon at the A site. Additionally, we find that *SufB2*, and likely all +1-frameshifting tRNAs, shifts to the +1-frame during the subsequent translocation reaction in which the translational GTPase elongation factor (EF)-G catalyzes the movement of *SufB2* from the A to P sites (i.e., a triplet-slippage mechanism). More specifically, we show that this frameshift occurs in the later steps of translocation, during which EF-G catalyzes a series of conformational rearrangements of the ribosomal pre-translocation (PRE) complex that enable the tRNA ASLs and their associated codons to move to their respective post-translocation positions within the ribosomal small (30S in bacteria) subunit^22–28^. Thus, efforts to increase the recoding efficiency of +1-frameshifting tRNAs should focus on enforcing a triplet anticodon-codon pairing in the 0-frame at the A site and directed evolution to optimize conformational rearrangements of the ribosomal PRE complex during the late stages of translocation.

**Figure 1.**
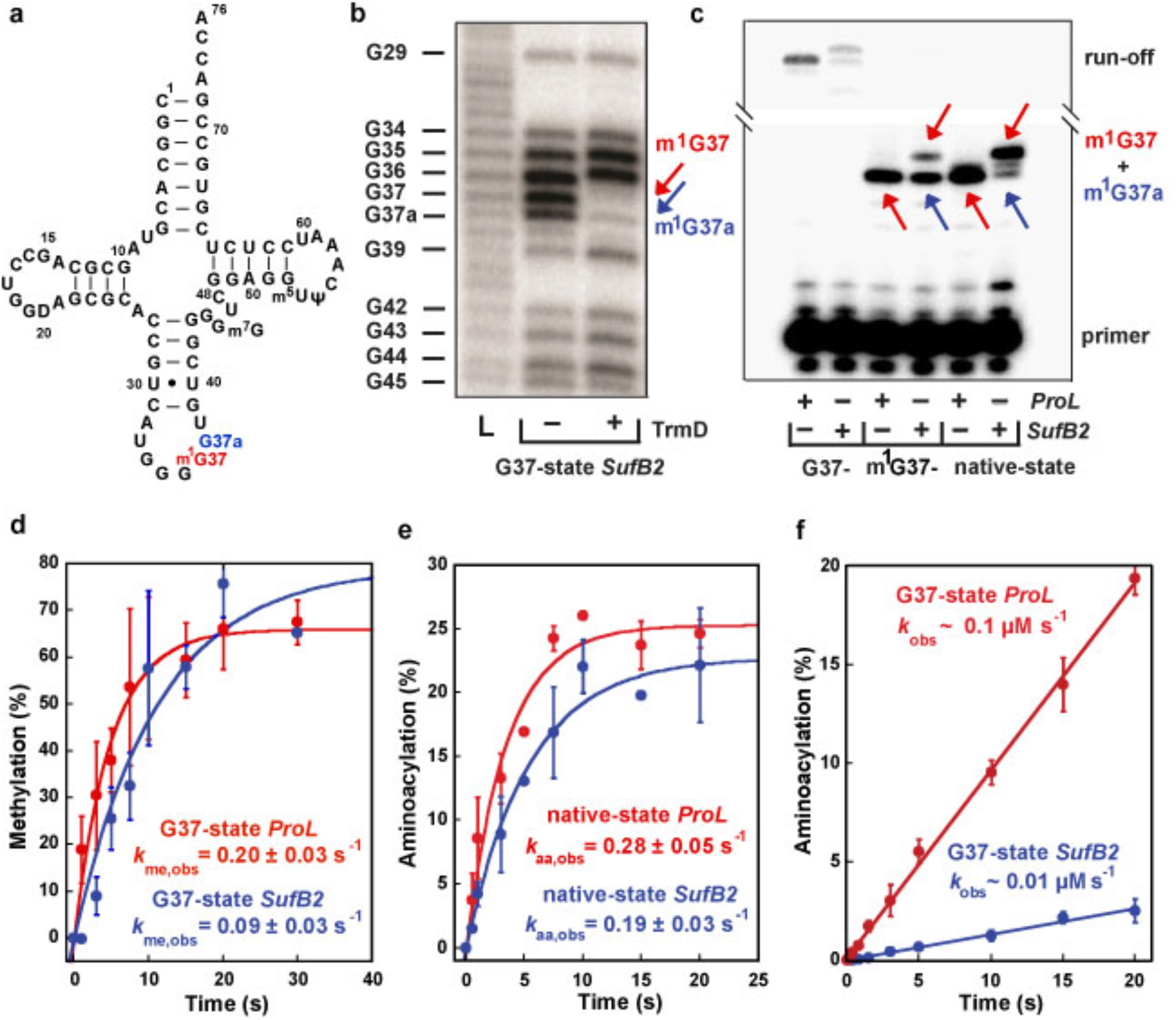
Methylation and aminoacylation of *SufB2* and *ProL*. **a** Sequence and secondary structure of native-state *SufB2*, showing the *N*^1^-methylated G37 in red and the G37a insertion to *ProL* in blue. **b** RNase T1 cleavage inhibition assays of TrmD-methylated G37-state *SufB2* transcript confirm the presence of m^1^G37 and m^1^G37a. Cleavage products are marked by the nucleotide positions of Gs. L: the molecular ladder of tRNA fragments generated from alkali hydrolysis. **c** Primer extension inhibition assays identify m^1^G37 in native-state *SufB2*. Red and blue arrows indicate positions of primer extension inhibition products at the methylated G37 and G37a, respectively, which are offset by one nucleotide relative to *ProL*. The first primer extension inhibition product for *SufB2* corresponds to m^1^G37a, the second corresponds to m^1^G37, while the primer extension inhibition product for *ProL* corresponds to m^1^G37. Due to the propensity of primer extension to make multiple stops on a long transcript of tRNA, the read-through primer extension product (54-55 nucleotides) had a reduced intensity relative to the primer extension inhibition products (21-22 nucleotides). Molecular size markers are provided by the primer alone (17 nucleotides) and the run-off products (54-55 nucleotides). **d** TrmD-catalyzed *N*^1^ methylation of G37-state *SufB2* and *ProL* as a function of time. **e, f** ProRS-catalyzed aminoacylation. **e** Aminoacylation of native-state *SufB2* and *ProL*. **f Aminoacylation of** G37-state *SufB2* and *ProL* as a function of time. In panels b, c, gels were performed three times with similar results, while in panels d-f, the bars are SD of three independent (n = 3) experiments, and the data are presented as mean values ± SD.

## RESULTS

### Native-state *SufB2* is *N*^1^-methylated at G37 and is readily aminoacylated with Pro

*SufB2* contains an extra G37a nucleotide inserted between G37 and U38 of *ProL*^20^ (Figure 1a). Whether the extra G37a is methylated and how it affects methylation of G37 is unknown. We thus determined the methylation status of the G37-G37a motif using RNase T1 cleavage inhibition assays and primer extension inhibition assays. We first generated a plasmid-encoded *SufB2* by inserting G37a into an existing Tac-inducible plasmid encoding *Escherichia coli ProL*^29^, which has an identical sequence to *S. typhimurium ProL.* We then expressed and purified the plasmid-encoded *SufB2* and *ProL* from an *E. coli ProL* knock-out (*ProL*-KO) strain^30^ containing all the endogenous enzymes necessary for processing *SufB2* and *ProL* to their *S. typhimurium* native states such that they possess the full complement of naturally occurring post-transcriptional modifications (termed the native-state tRNAs). In addition, we prepared in vitro transcripts of *SufB2* and *ProL* lacking all post-transcriptional modifications (termed the G37-state tRNAs), or enzymatically methylated with purified *E. coli* TrmD^30,31^ such that they possess only the *N*^1^-methylation at G37 and no other post-transcriptional modifications (termed the m^1^G37-state tRNAs). In the case of *SufB2*, RNase T1 cleavage inhibition assays demonstrated cleavage at G37 and G37a of the G37-state tRNA, but inhibition of cleavage at either position upon treatment with TrmD (Figure 1b), indicating that both nucleotides are *N*^1^-methylated in the m^1^G37-state tRNA.

Primer extension inhibition assays, which were previously validated by mass spectrometry analysis^30^, showed inhibition of extension at G37 and G37a in m^1^G37- and native-state *SufB2* (Figure 1c), confirming that both nucleotides are *N*^1^-methylated in these species. Notably, *N*^1^ methylation shifted almost entirely to G37 in native-state *SufB2*, indicating that m^1^G37 is the dominant methylation product in cells. As a control, no inhibition of extension at G37 or G37a was observed for G37-state *SufB2.* Complementary kinetics experiments showed that the yield and rate of *N*^1^-methylation of G37-state *SufB2* were similar to those of G37-state *ProL* (Figure 1d). Likewise, kinetics experiments revealed that the yield and rate of aminoacylation of native-state *SufB2* with Pro were similar to those of native-state *ProL* (Figure 1e). In contrast, aminoacylation of G37-state *SufB2* was inhibited (Figure 1f). These results demonstrate that the native-state *SufB2* synthesized in cells is quantitatively *N*^1^-methylated to generate m^1^G37 and is readily aminoacylated with Pro.

### *SufB2* promotes +1 frameshifting using triplet-slippage and possibly other mechanisms

We next determined the mechanism(s) through which *SufB2* promotes +1 frameshifting in a cellular context. We created a pair of isogenic *E. coli* strains expressing *SufB2* or *ProL* from the chromosome in a *trmD-knockdown (trmD-KD)* background^30^. This background strain was designed to evaluate the effect of m^1^G37 on +1 frameshifting and it was generated by deleting chromosomal *trmD* and controlling cellular levels of m^1^G37 using arabinose-induced expression of the human counterpart *trm5*, which is competent to stoichiometrically *N*^1^-methylate intracellular tRNA substrates^30^. The isogenic pair of the *SufB2* and *ProL* strains were measured for +1 frameshifting in a cell-based *lacZ* reporter assay in which a CCC-C motif was inserted into the 2^nd^ codon position of *lacZ* such that a +1-frameshifting event at the motif was necessary to synthesize full-length β-galactosidase (β-Gal)^29^. The efficiency of +1 frameshifting was calculated as the ratio of β-Gal expressed in cells containing the CCC-C insertion relative to cells containing an in-frame CCC insertion.

In the m^1^G37-abundant (m^1^G37+) condition, *SufB2* displayed a high +1-frameshifting efficiency (8.2%, Figure 2a) relative to *ProL* (1.4%). In the m^1^G37-deficient (m^1^G37-) condition, *SufB2* exhibited an even higher efficiency (20.8%) and, consistent with our previous work^29^, *ProL* also displayed an increased efficiency (7.0%) relative to background (1.4%). Because *N*^1^-methylation in the m^1^G37+ condition was stoichiometric (Figure 1c), thereby preventing quadruplet-pairing, we attribute the 8.2% efficiency of *SufB2* in this condition as arising exclusively from triplet-slippage. In the m^1^G37-condition, we observed an increase in +1-frameshifting efficiency of *SufB2* to 20.8%. While multiple mechanisms may exist for the increased +1 frameshifting, the exploration of both triplet-slippage and quadruplet-pairing is one possibility.

**Figure 2.**
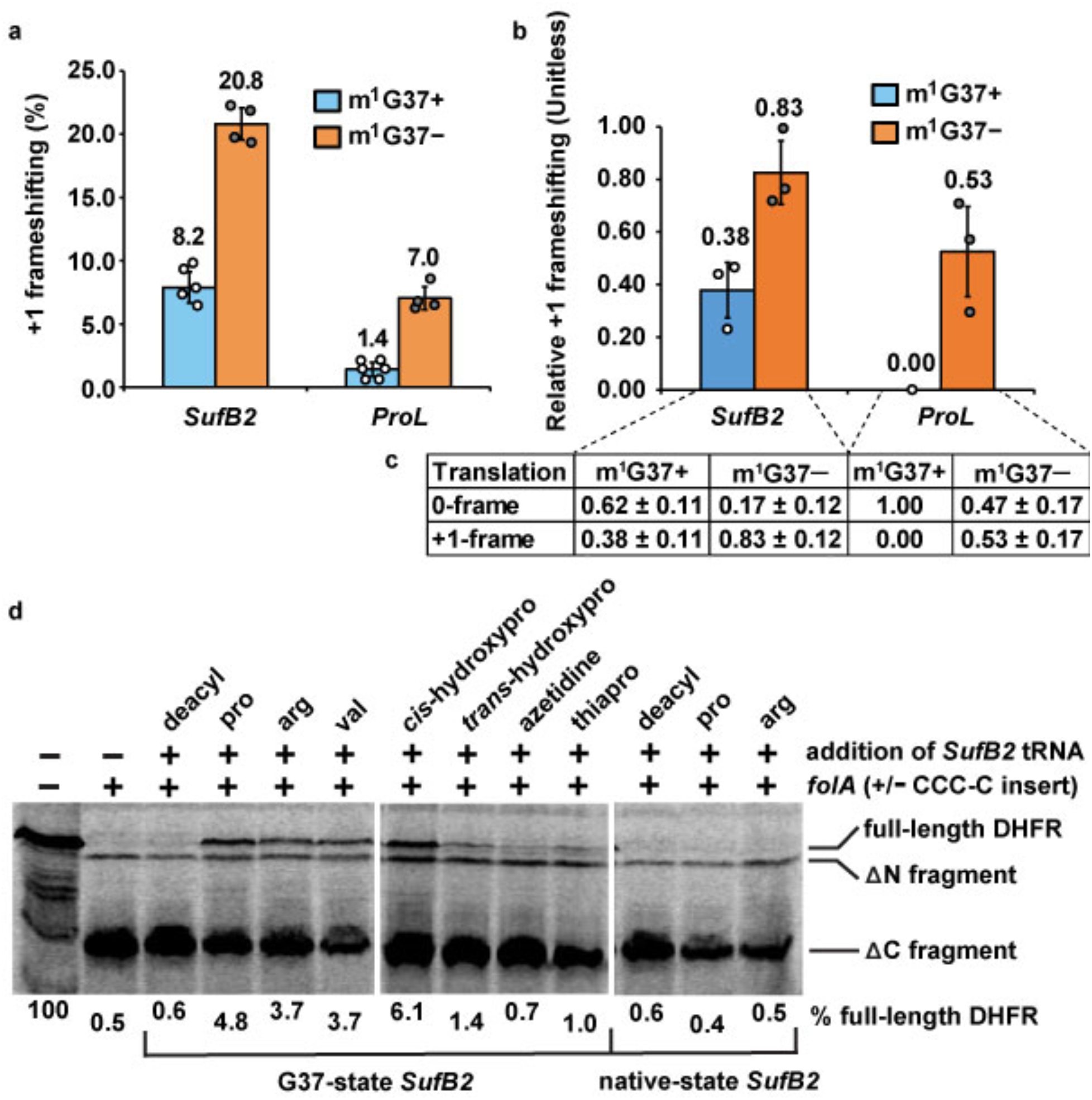
*SufB2*-induced +1 frameshifting and genome recoding. **a** The +1-frameshifting efficiency in cell-based *lacZ* assay for *SufB2* and *ProL* strains in m^1^G37+ and m^1^G37-conditions. The bars in the graph are SD of four, five, or six independent (n = 4, 5, or 6) biological repeats, and the data are mean values ± SD. **b** The difference in the ratio of protein synthesis of *lolB* to *cysS* for *SufB2* and *ProL* strains in m^1^G37+ and m^1^G37-conditions relative to *ProL* in the m^1^G37+ condition. **c** Measurements underlying the bar plots in panel **b**. Each ratio was measured directly and the ratio of *ProL* in the m^1^G37+ condition was normalized to 1.0. The difference of each ratio relative to the normalized ratio represented the +1-frameshifting efficiency at the CCC-C motif at the 2^nd^ codon of *lolB*. The bars in the graph are SD of three independent (n = 3) biological repeats, and the data are mean values ± SD. In **a, b**, decoding of the CCC-C motif was mediated by *SufB2* and *ProM* in the *SufB2* strain, and by *ProL* and *ProM* in the *ProL* strain, where the presence of *ProM* ensured no vacancy at the CCC-C motif. The increased +1 frameshifting in the m^1^G37-condition vs. the m^1^G37+ condition indicates that *SufB2* and *ProL* are each an active determinant in decoding the CCC-C motif. **d** *SufB2*-mediated insertion of non-proteinogenic amino acids at the CCC-C motif in the 5^th^ codon position of *folA* using [^35^S]-Met-dependent in vitro translation. Reporters of *folA* are denoted by +/- CCC-C, where “+” and “-” indicate constructs with and without the CCC-C motif. SDS-PAGE analysis identifies full-length DHFR resulting from a +1-frameshift event at the CCC-C motif by *SufB2* pre-aminoacylated with the amino acid shown at the top of each lane, a DC fragment resulting from lack of the +1-frameshift event, and a ΔN fragment resulting from translation initiation at the AUG codon likely at position 17 or 21 downstream from the CCC-C motif. Gel samples were derived from the same experiment, which was performed five times with similar results. Gels for each experiment were processed in parallel. Lane 1: full-length DHFR as the molecular marker; deacyl: deacylated tRNA.

To confirm our results, we performed similar studies with the isogenic *SufB2* and *ProL* strains on the endogenous *E. coli lolB* gene, encoding the outer membrane lipoprotein. The *lolB* gene naturally contains a CCC-C motif at the 2^nd^ codon position such that +1 frameshifting at this motif would decrease protein synthesis due to premature termination. As a reference, we used *E. coli cysS*, encoding cysteinyl-tRNA synthetase (CysRS)^30^, which has no CCC-C motif in the first 16 codons and would be less sensitive to +1 frameshifting at CCC-C motifs during protein synthesis. The ratio of protein synthesis of *lolB* to *cysS* for the control sample *ProL* in the m^1^G37 condition, measured from Western blots (Methods), was normalized to 1.00, denoting that *lolB* and *cysS* were maximally translated in the 0-frame without +1 frameshifting (i.e., a relative +1 frameshifting efficiency of 0.00) (Figures 2b, 2c). In the m^1^G37+ condition, *SufB2* displayed a ratio of LolB to CysRS of 0.62, indicating an increase in the relative +1 frameshifting efficiency to 0.38, and in the m^1^G37–condition, it displayed a ratio of 0.17, indicating an increase in the relative +1 frameshifting efficiency to 0.83 (Figures 2b, 2c). Similarly, *ProL* in the m^1^G37–condition displayed a ratio of LolB to CysRS of 0.47, indicating an increase in the +1-frameshifting efficiency to 0.53.

### *SufB2* can insert non-proteinogenic amino acids at CCC-C motifs

We next asked whether *SufB2* can deliver non-proteinogenic amino acids to the ribosome by inducing +1 frameshifting at a CCC-C motif (Figure 2d). We inserted a CCC-C motif at the 5^th^ codon position of the *E. coli folA* gene, encoding dihydrofolate reductase (DHFR). A *SufB2*-induced +1 frameshifting event at the insertion would result in full-length DHFR, whereas the absence of +1 frameshifting would result in a C-terminal truncated DHFR fragment (DC). *SufB2* was aminoacylated with non-proteinogenic amino acids using a Flexizyme^32^ and subsequently tested in [^35^S]-Met-dependent in vitro translation reactions using the *E. coli* PURExpress system. The resulting protein products were separated by sodium dodecyl sulfate (SDS)-polyacrylamide gel electrophoresis and quantified by phosphorimaging. Control experiments with no *SufB2* or with a non-acylated *SufB2* showed no full-length DHFR, demonstrating that synthesis of full-length DHFR depended upon *SufB2* delivery of an amino acid as a result of +1 frameshifting at the CCC-C motif. We showed that *SufB2* was able to deliver Pro, Arg, Val, and the Pro analogs *cis*-hydroxypro, *trans*-hydroxypro, azetidine, and thiapro (Supplementary Figure 1) to the ribosome in response to the CCC-C motif, and that the efficiency of delivery by G37-state *SufB2* was generally higher than that by native-state *SufB2*. Notably, the PURExpress system contains all canonical tRNAs, including *ProL* and *ProM*, indicating the ability of *SufB2* to successfully compete with these tRNAs.

### *SufB2* uses triplet pairing in the 0-frame at the A site

To determine at which step in the elongation cycle *SufB2* undergoes +1 frameshifting in response to a CCC-C motif, we used an *E. coli* in vitro translation system composed of purified components and supplemented with requisite tRNAs and translation factors to perform a series of ensemble rapid kinetic studies. We began with a GTPase assay that reports on the yield and rate with which the translational GTPase EF-Tu hydrolyzes GTP upon delivery of a ternary complex (TC), composed of EF-Tu, [γ-^32^P]-GTP, and prolyl-*SufB2* (*SufB2*-TC) or *ProL (ProL-TC)*, to the A site of a ribosomal 70S initiation complex (70S IC) carrying an initiator fMet-tRNA^fMet^ in the P site and a programmed CCC-C motif at the A site. The results of these experiments showed that the yield and rate of GTP hydrolysis (*k*_GTP,obs_) upon delivery of *SufB2-TC* were quantitatively similar to those of *ProL*-TC for both the native- and G37-state tRNAs (Figure 3a).

**Figure 3.**
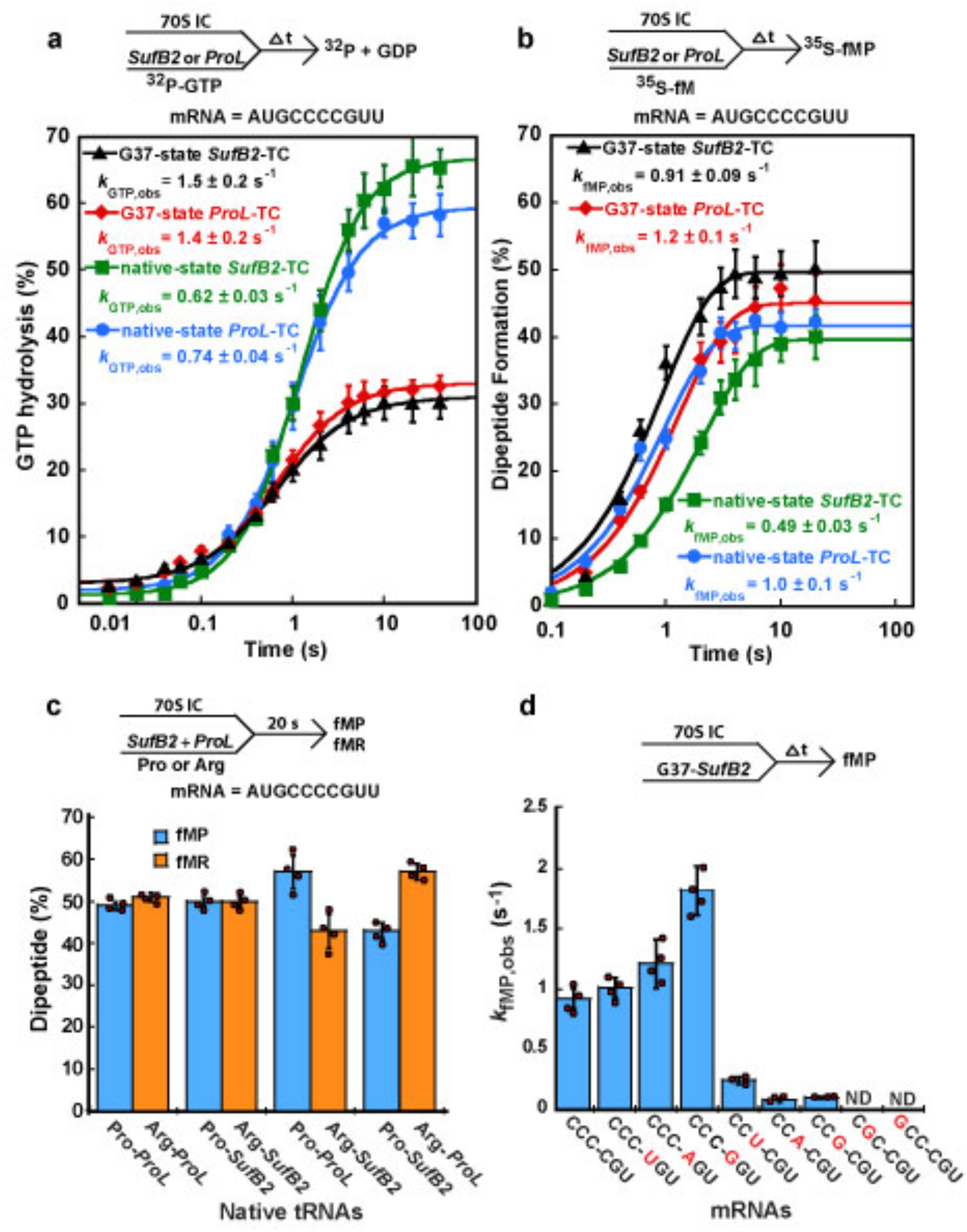
*SufB2* uses a triplet anticodon-codon pairing scheme at the A site. **a** GTP hydrolysis by EF-Tu as a function of time for delivery of G37- or native-state *SufB2-* or *ProL*-TC to the A site of a 70S IC. Although the concentration of TCs was limiting, which would limit the rate of binding of TCs to the 70S IC, the observed differences in the yield of GTPase activity indicated that binding was not the sole determinant, but that other factors, such as the identity and the methylation state of the tRNA, affected the GTPase activity. **b** Dipeptide fMP formation as a function of time for delivery of G37- or native-state *SufB2-* or *ProL*-TC to the A site of a 70S IC. Due to the limiting concentration of the 70S IC, which did not include the tRNA substrate, the yield of di-or tri-peptide formation assays was constant even with different tRNAs in TCs. **c** The yield of fMP and fMR in dipeptide formation assays in which equimolar mixtures of native-state *SufB2*-TC, carrying Pro and/or Arg, and/or native-state *ProL*-TC, carrying Pro and/or Arg, are delivered to 70S ICs. The mRNA in 70S ICs in (A-C) is AUG-CCC-CGU-U. **d** Dipeptide formation rate *k*_fMP,obs_ for delivery of G37-state *SufB2*-TC to 70S ICs containing sequence variants of the CCC-C motif in the A site. In panels **a, b**, the bars in the graphs are SD of three independent (n = 3) experiments, in panel **c**, the bars in the graphs are SD of four independent (n = 4) experiments, and in panel **d**, the bars in the graphs are SD of three or four independent (n = 3 or 4) experiments. All data are presented as mean values ± SD. Δt: a time interval, ND: not detected.

We next performed a dipeptide formation assay that reports on the synthesis of a peptide bond between the [^35^S]-fMet moiety of a P-site [^35^S]-fMet-tRNA^fMet^ in a 70S IC and the Pro moiety of a *SufB2-* or *ProL*-TC delivered to the A site. This assay revealed that the rate of [^35^S]-fMet-Pro (fMP) formation (*k*_fMP,obs_) for *SufB2-TC* was within 2-fold of that for *ProL*-TC for both the native- and G37-state tRNAs (Figure 3b, Table S2).

To test whether native-state *SufB2*-TC can effectively compete with *ProL*-TC for delivery to the A site and peptide-bond formation, we varied the dipeptide formation assay such that an equimolar mixture of each TC was used in the reaction (Figure 3c). Since aminoacylation of both tRNAs with Pro would create dipeptides of the same identity (i.e., fMP), we used a Flexizyme to aminoacylate them with different amino acids and generate distinct dipeptides. Control experiments showed that *ProL* charged with Pro or Arg (Figure 3c, Bars 1 and 2) and *SufB2* charged with Pro or Arg (Bars 3 and 4) generated the same amount of fMP and fMR, indicating that the amino-acid identity did not affect the level of dipeptide formation. We found that the amount of dipeptide formed by *SufB2-TC* and *ProL-TC* in these competition assays was similar, although the amount formed by *SufB2*-TC was slightly less (45% vs. 55%), in both the native- (Bars 5-8) and G37-state tRNAs (Supplementary Figure 2a). These competition experiments provide direct evidence that *SufB2*-TC effectively competes with *ProL-TC* for delivery to the A site and peptide-bond formation.

Collectively, the results of our GTPase-, dipeptide formation-, and competition assays indicate that *SufB2*-TC is delivered to the A site and participates in peptide-bond formation in the same way as *ProL*-TC, suggesting that *SufB2* uses triplet pairing in the 0-frame at the A site that successfully competes with triplet pairing by *ProL*. To support this interpretation, we measured *k*_fMP,obs_ in our dipeptide formation assay, using G37-state *SufB2*-TC and a series of mRNA variants in which single nucleotides in the CCC-C motif were substituted. We showed that *K*_fMP,obs_ did not decrease upon substitution of the 4^th^ nucleotide of the CCC-C motif, but that it decreased substantially upon substitution of any of the first three nucleotides of the motif (Figure 3d, Supplementary Figure 2b). Thus, triplet pairing of *SufB2* to the first three Cs of the CCC-C motif is necessary and sufficient for rapid delivery of the tRNA to the A site and its participation in peptide-bond formation.

### The A-site activity of *SufB2* depends on the sequence of the anticodon loop

We next asked how delivery of *SufB2-TC* to the A site and peptide-bond formation depend on the sequence of the *SufB2* anticodon loop. Starting from G37-state *SufB2*, we created two variants containing a G-to-C substitution in nucleotide 37 (G37C) or 34 (G34C) within the anticodon loop and adapted our dipeptide formation assay to measure the fMP yield and *K*_fMP,obs_ generated by each variant at the CCC-C motif at the A site. We showed that the G37C variant resulted in a fMP yield of 32% and a *K*_fMP,obs_ of 0.14 ± 0.01 s^-1^, most likely by triplet pairing of nucleotides 34-36 of the anticodon loop with the 0-frame of the CCC-C motif (Figure 4a). In contrast, the G34C variant resulted in a fMP yield of 30% and a *k*_fMP,obs_ of 0.28 ± 0.04 s^-1^, most likely by triplet pairing of nucleotides 35-37 of the anticodon loop with the 0-frame of the CCC-C motif (Figure 4b). Our interpretation that nucleotides 35-37 of the anticodon loop of the G34C variant most likely triplet pair with the 0-frame of the CCC-C motif is consistent with the observations that the fMP yield and *k*_fMP,obs_ of the G34C variant are similar and 2-fold higher, respectively, than those of the G37C variant. If nucleotides 34-36 of the anticodon loop of the G34C variant were to form a triplet pair with the CCC-C motif, we would have expected it to pair in the +2-frame, which would have most likely reduced the fMP yield and *k*_fMP,obs_ of the G34C variant relative to the G37C variant. These results suggest that G37-state *SufB2* exhibits some plasticity as to whether it can undergo triplet pairing with anticodon loop nucleotides 34-36 or 35-37, consistent with a previous study^33^.

**Figure 4.**
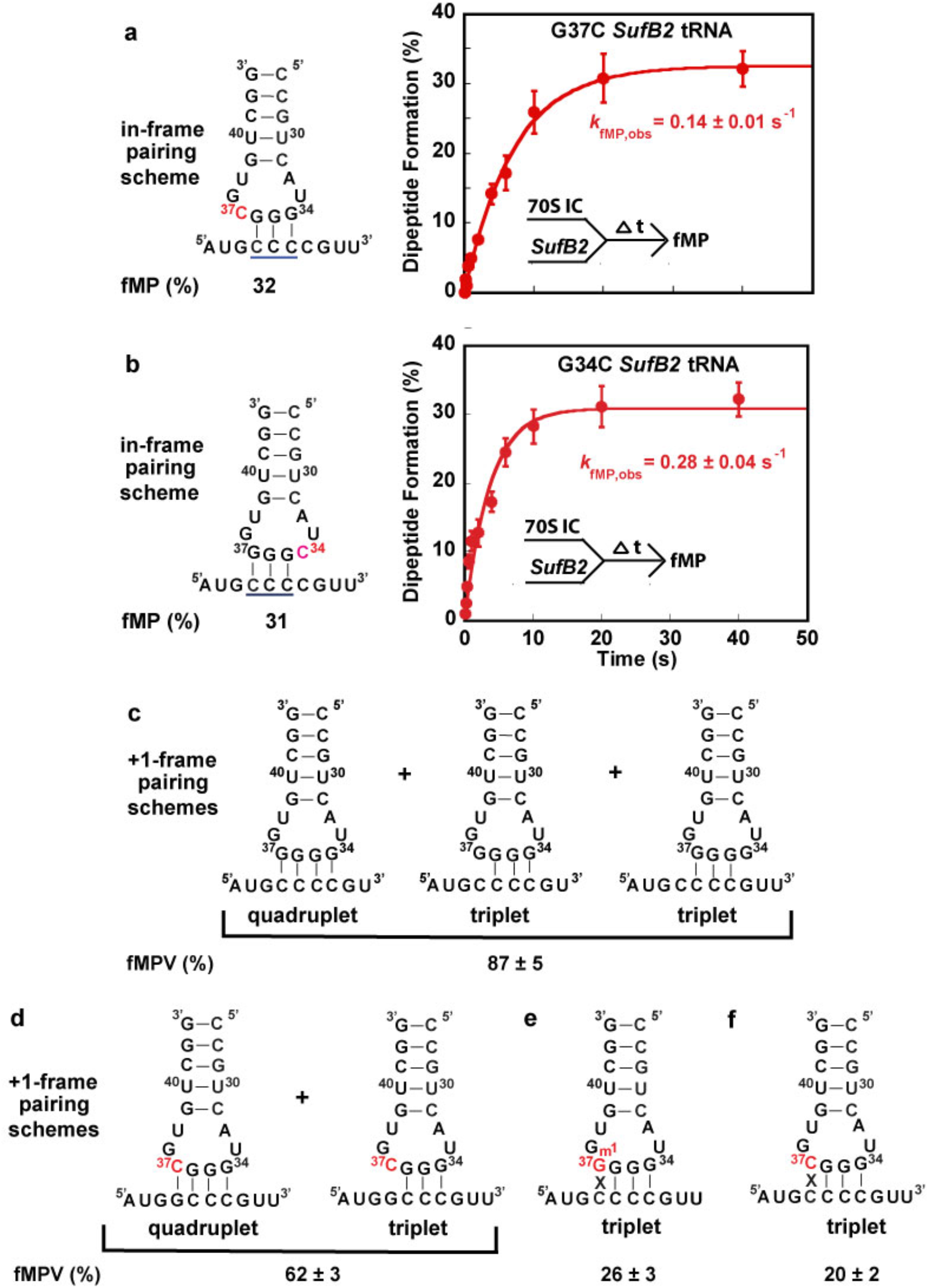
Plasticity of *SufB2*-induced +1 frameshifting. **a** fMP formation as a function of time upon delivery of the G37C variant of G37-state *SufB2*-TC to the A site of a 70S IC, allowing nucleotides 34-36 to pair with a CCC-C motif at the A site. **b** fMP formation as a function of time upon delivery of the G34C variant of G37-state *SufB2*-TC to the A site of a 70S IC, allowing nucleotides 35-37 to pair with a CCC-C motif. **c-f** Results of fMPV formation assays in which *SufB2*-TC is delivered to an A site programmed with a quadruplet codon at the 2^nd^ position and sequences of the *SufB2* anticodon loop and/or quadruplet codon are varied. Yields of fMPV formation represent +1 frameshifting during translocation of *SufB2* from the A site to the P site. Possible +1-frame anticodon-codon pairing schemes of *SufB2* during translocation: **c** G37-state *SufB2* capable of frameshifting at a CCC-C motif via quadruplet pairing and/or triplet slippage, **d** G37C variant of G37-state *SufB2* capable of frameshifting at a GCC-C motif via quadruplet pairing and/or triplet slippage, **e** m^1^G37-state *SufB2* capable of frameshifting at a CCC-C motif via only triplet slippage, and **f** G37C variant of G37-state *SufB2* capable of frameshifting at a CCC-C motif via only triplet slippage. In panels **a, b**, the bars in the graphs are SD of three (n = 3) independent experiments, and the data are presented as mean values ± SD. Δt: a time interval.

### *SufB2* shifts to the +1-frame during translocation

Although *SufB2* uses triplet pairing in the 0-frame when it is delivered to the A site, it is a highly efficient +1-frameshifting tRNA (Figure 2). We therefore asked whether +1 frameshifting occurs during or after translocation of *SufB2* into the P site. We addressed this question by adapting our previously developed tripeptide formation assays^29^. We rapidly delivered EF-G and an equimolar mixture of G37-state *SufB2*-, tRNA^Val^-, and tRNA^Arg^-TCs to 70S ICs assembled on an mRNA in which the 2^nd^ codon was a CCC-C motif and the 3^rd^ codon was either a GUU codon encoding Val in the +1 frame or a CGU codon encoding Arg in the 0-frame. As soon as translocation of the PRE complex and the associated movement of SufB2 from the P to A sites formed a ribosomal post-translocation (POST) complex with an empty A site in these experiments, tRNA^Val^- and tRNA^Arg^-TC would compete for the codon at the A site to promote formation of an fMPV tripeptide or an fMPR tripeptide. Thus, the fMPV yield and *k*_fMPV,obs_ report on the sub-population of *SufB2* that shifted to the +1-frame, whereas the fMPR yield and *k*_fMPR,obs_ report on the sub-population that remained in the 0-frame^29,34^. The results showed that the yield of fMPV was much higher than that of fMPR (90% vs. 10%, Figure 5a), demonstrating the high efficiency with which G37-state *SufB2* induces +1 frameshifting. Notably, relative to the +1 frameshifting of *ProL* we have previously reported^29^, *K*_fMPV,obs_ of *SufB2* (0.09 s^-1^) was comparable to the rate of +1 frameshifting of *ProL* during translocation (0.1 s^-1^) rather than that of +1 frameshifting after translocation into the P site (~10^-3^ s^-1^)^29^, indicating that *SufB2* underwent +1 frameshifting during translocation. Our observation that the fMPV yield plateaus at 90% at long reaction times suggests that the sub-populations of *SufB2* that will shift to the +1-frame and remain in the 0-frame are likely established in the A site, even before EF-G binds to the PRE complex. Given that *SufB2* exhibits triplet pairing in the 0-frame at the A site (Figures 3a-c, Supplementary Table 2, and Supplementary Figure 2a) and shifts into the +1-frame during translocation (Figure 5a), the two sub-populations of *SufB2* in the A site seem to differ primarily in their propensity to undergo +1 frameshifting during translocation. The sub-population that encompasses 90% of the total would exhibit a high propensity of undergoing +1 frameshifting during translocation, whereas the sub-population that encompasses 10% of the total would exhibit a low propensity of undergoing +1 frameshifting during translocation, preferring instead to remain in the 0-frame.

**Figure 5.**
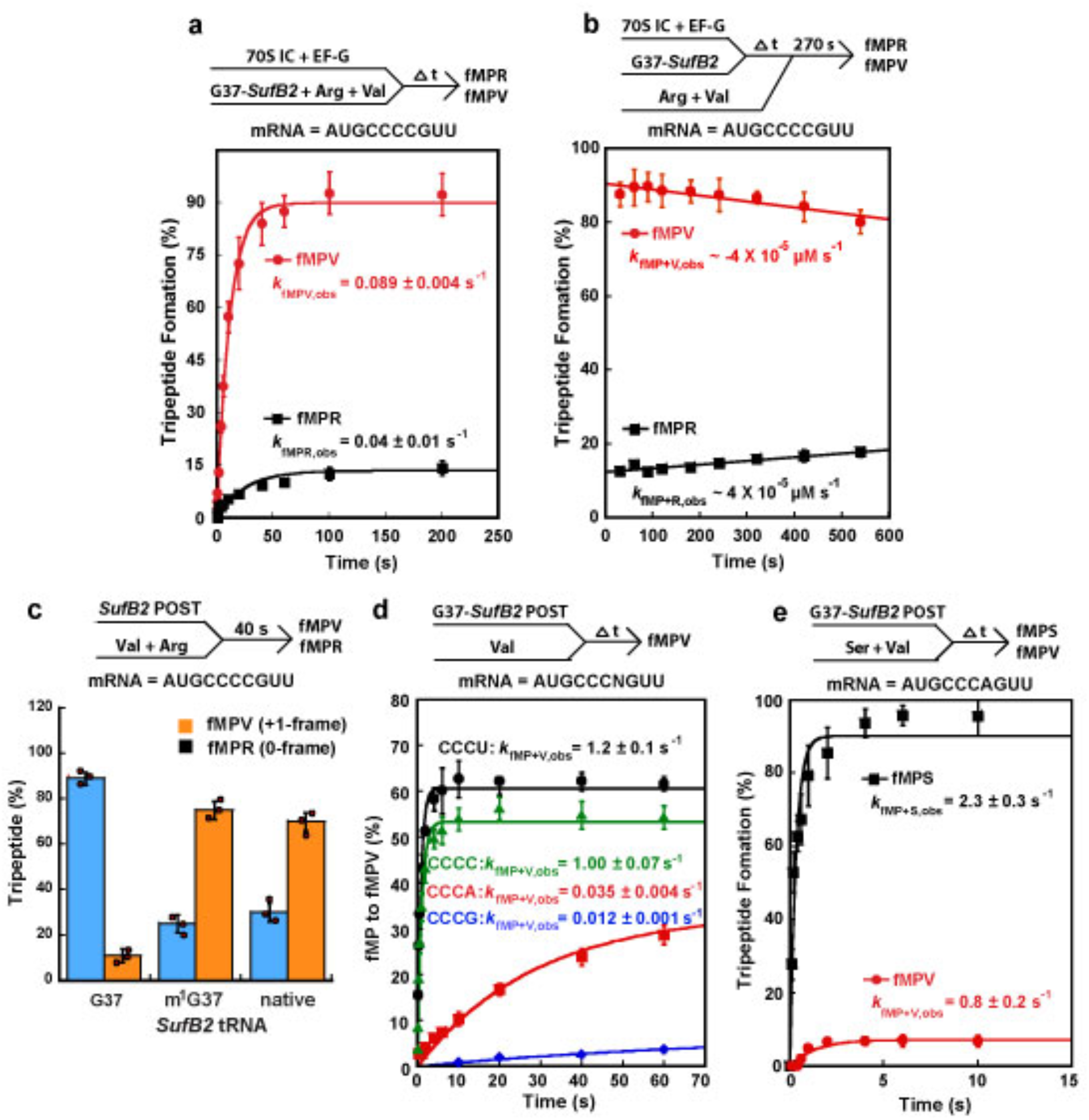
*SufB2* shifts to the +1-frame during translocation. **a** Relative fMPV and fMPR formation as a function of time upon rapid delivery of EF-G and an equimolar mixture of G37-state *SufB2-*, tRNA^Val^-, and tRNA^Arg^-TCs to 70S ICs carrying a CCC-C motif in the A site. **b** Relative fMPV and fMPR formation as a function of time when a defined time interval is introduced between delivery of G37-state *SufB2*-TC and EF-G and delivery of an equimolar mixture of tRNA^Arg^- and tRNA^Val^-TCs. **c** Relative fMPV and fMPR formation after reacting fMP-POST complexes with a mixture of tRNA^Val^- and tRNA^Arg^-TCs based on the time courses in Supplementary Figures 2**d-f**. **d** fMPV formation as a function of time upon rapid delivery of tRNA^Val^-TC to an fMP-POST complex carrying a CCC-N motif in the A site. **e** Relative fMPV and fMPS formation as a function of time upon rapid delivery of an equimolar mixture of tRNA^Val^- and tRNA^Ser^-TCs to an fMP-POST complex carrying a CCC-A motif in the A site. In panels **a-e**, the bars are SD of three (n = 3) independent experiments and the data are presented as mean values ± SD. Arg: arginyl-tRNA^Arg^; Val: valyl-tRNA^Val^.

We next determined whether the 10% sub-population of G37-state *SufB2* that remained in the 0-frame during translocation could undergo +1 frameshifting after arrival at the P site. We varied our tripeptide formation assay so as to deliver the TCs in two steps separated by a defined time interval (Figure 5b). In the first step, G37-state *SufB2*-TC and EF-G were delivered to the 70S IC to form a POST complex, which was then allowed the opportunity to shift to the +1-frame over a systematically increasing time interval. In the second step, an equimolar mixture of tRNA^Arg^- and tRNA^Val^-TCs was delivered to the POST complex. The results showed that fMPV was rapidly formed at a high yield and exhibited a *K*_fMP+V,obs_ (where the “+” denotes the time interval between the delivery of translation components) that did not increase as a function of time. In contrast, fMPR was formed at a low yield and exhibited a *K*_fMP+R,obs_ that did not decrease as a function of time. Together, these results indicate that the sub-population of P site-bound *SufB2* in the 0-frame does not undergo +1 frameshifting. This interpretation is supported by the observation that EF-P, an elongation factor which we showed suppresses +1 frameshifting within the P site^29^, had no effect on the yield of fMPV yield (Supplementary Figure 2c and Supplementary Table 3).

Having shown that +1 frameshifting of *SufB2* occurs only during translocation, we evaluated the effect of m^1^G37 on the frequency of this event. We began by delivering G37-, m^1^G37-, or native-state *SufB2-TCs* together with EF-G to 70S ICs to form the corresponding POST complexes and then delivered an equimolar mixture of tRNA^Arg^- and tRNA^Val^-TCs to each POST complex to determine the relative formation of fMPV and fMPR. The results showed that m^1^G37- and native-state *SufB2* displayed a reduced fMPV yield and a concomitantly increased fMPR yield relative to G37-state *SufB2* (Figures 5c, Supplementary Figures 2d-f), consistent with the notion that the presence of m^1^G37 compromises +1 frameshifting.

We then used the same tripeptide formation assay to determine how +1 frameshifting during translocation of G37-state *SufB2* depends on the identity of the 4^th^ nucleotide of the CCC-C motif. A series of POST complexes were generated by delivering G37-state *SufB2-TCs* and EF-G to 70S ICs programmed with a CCC-N motif at the 2^nd^ codon position. Each POST complex was then rapidly mixed with tRNA^Val^-TC to monitor the yield of fMPV and *k*_fMP+V,obs_ (Figure 5d). The results showed a high fMPV yield and high *k*_fMP+V,obs_ at the CCC-[C/U] motifs, but a low yield and low *k*_fMP+V,obs_ at the CCC-[A/G] motifs. This indicates that high-efficiency of *SufB2*-induced +1 frameshifting during translocation requires the presence of a [C/U] at the 4^th^ nucleotide of the CCC-C motif. Because *SufB2* in these experiments was in the G37-state, it is possible that a sub-population underwent +1 frameshifting via quadruplet-pairing with the [C/U] at the 4^th^ nucleotide of the CCC-[C/U] motif during translocation. It is also possible that a sub-population underwent +1 frameshifting via triplet-slippage, which could potentially be inhibited by the presence of [G/A] at the 4^th^ nucleotide of the motif. To verify that the POST complex formed with the CCC-A sequence was largely in the 0-frame, we rapidly mixed the complex with an equimolar mixture of tRNA^Ser^-TC, cognate to the next A-site codon in the 0-frame (AGU), and tRNA^Val^-TC, cognate to the next A-site codon in the +1-frame (GUU) (Figure 5e). The results showed a high yield and high *k*_fMP+S,obs_, supporting the notion that the POST complex formed with the CCC-A motif was largely in the 0-frame. Thus, the 4^th^ nucleotide of the CCC-C motif plays a role in determining +1 frameshifting during translocation of *SufB2* from the A site to the P site.

### The +1-frameshifting efficiency of *SufB2* depends on sequences of the anticodon loop and the CCC-C motif

To determine whether the +1-frameshifting efficiency of *SufB2* during translocation is influenced by sequences of the anticodon loop and the CCC-C motif, we performed tripeptide formation assays and monitored the yield of fMPV. In these experiments, we varied the sequence of the *SufB2* anticodon loop and/or the CCC-C motif at the 2^nd^ codon position of the mRNA. To explore the possibilities of both triplet-slippage and quadruplet-pairing, we used variants of G37-state *SufB2*. We showed that variants with the potential to undergo quadruplet-pairing with the CCC-C motif resulted in fMPV yields of 87% and 62% (Figures 4c, d). The different yields suggest that G37-state *SufB2* variants can induce triplet-slippage and/or engage in quadruplet-pairing with different efficiencies during translocation. Analogous experiments showed that *SufB2* variants that were restricted to triplet-pairing resulted in reduced fMPV yields (26% and 20%, respectively) upon pairing with a CCC-C motif (Figures 4e, f). Collectively, these results suggest that there is considerable plasticity in the mechanisms that *SufB2* uses to induce +1 frameshifting during translocation and in the efficiencies of these mechanisms.

### An smFRET signal that reports on ribosome dynamics during individual elongation cycles

To address the mechanism of *SufB2*-induced +1 frameshifting during translocation, we used a previously developed smFRET signal to determine whether and how *SufB2* alters the rates with which the ribosome undergoes a series of conformational changes that drive and regulate the elongation cycle^35^ (Figures 6a-c). This signal is generated using a ribosomal large, or 50S, subunit that has been Cy3- and Cy5-labeled at ribosomal proteins bL9 and uL1, respectively, to report on ‘opening’ and ‘closing’ of the L1 stalk of the 50S subunit. Accordingly, individual FRET efficiency (*E*_FRET_) vs. time trajectories recorded using this signal exhibit transitions between two FRET states corresponding to the ‘open’ (*E*_FRET_ = ~0.55) and ‘closed’ (*E*_FRET_ = ~0.31) conformations of the L1 stalk (Figure 6d).

**Figure 6.**
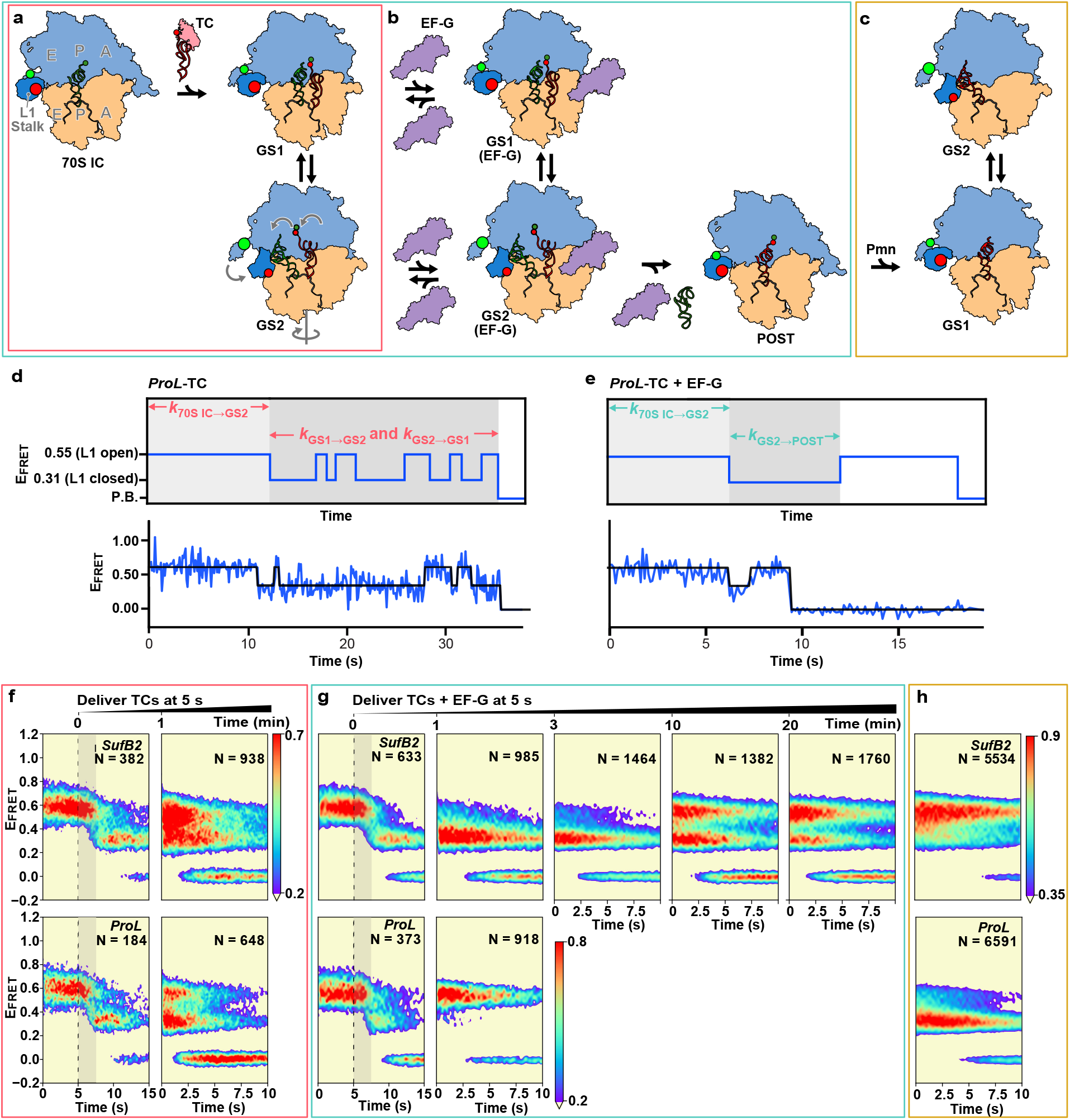
*SufB2* interferes with elongation complex dynamics during late steps of translocation. **a-c** Cartoon representation of elongation as a G37-state *SufB2-* or *ProL*-TC is delivered to the A site of a bL9(Cy3)- and uL1(Cy5)-labeled 70S IC; **a** in the absence, or **b** in the presence of EF-G, or **c** upon using puromycin (Pmn) to deacylate the P site-bound G37-state *SufB2* or *ProL* and generate the corresponding PRE^-A^ complex. The 30S and 50S subunits are tan and light blue, respectively; the L1 stalk is dark blue; Cy3 and Cy5 are bright green and red spheres, respectively; EF-Tu is pink; EF-G is purple; fMet-tRNA^fMet^ is dark green; and *SufB2* or *ProL* is dark red. **d, e** Hypothetical (top) and representative experimentally observed (bottom) *E*_FRET_ vs. time trajectories recorded as *ProL*-TC is delivered to a 70S IC, **d** in the absence and e in the presence of EF-G as depicted in **a, b**. The waiting times associated with *k*_70S IC→GS2_, *k*_GS1→GS2_, *k*_GS2→GS1_, and *k*_GS2→POST_ are indicated in each hypothetical trajectory. **f, g, and h** Surface contour plots of the time evolution of population FRET obtained by superimposing individual *E*_FRET_ vs. time trajectories in the experiments in **a**, **b**, and **c**, respectively, for *SufB2* (top) and *ProL* (bottom). N: the number of trajectories used to construct each contour plot. Surface contours are colored as denoted in the population color bars. For pre-steady-state experiments, the black dashed lines indicate the time at which the TC was delivered and the gray shaded areas denote the time required for the majority (54 - 68%) of the 70S ICs to transition to GS2. Note that the rate of deacylated *SufB2* dissociation from the A site under our conditions is similar to that of EF-G-catalyzed translocation, thereby resulting in the buildup of a PRE complex sub-population over 3-20 min post-delivery that lacks an A site tRNA and is incapable of translocation. This sub-population exhibits *k*_GS1→GS2_, *k*_GS2→GS1_, and *K*_eq_ values similar to those observed in experiments recorded in the absence of EF-G (Supplementary Table 6).

Previously, we have shown that open→closed and closed→open L1 stalk transitions correlate with a complex series of conformational changes that take place during an elongation cycle^35–37^. The L1 stalk initially occupies the open conformation as an aa-tRNA is delivered to the A site of a 70S IC or POST complex and peptide-bond formation generates a PRE complex that is in a global conformation we refer to as global state (GS) 1. The PRE complex then undergoes a large-scale structural rearrangement that includes an open→closed transition of the L1 stalk so as to occupy a second global conformation we refer to as GS2 (i.e., the 0.55→0.31 *E*_FRET_ transition denoted by the rate *k*_70S IC→GS2_ in Figures 6d and e, corresponding to the multi-step 70S IC→GS2 transition in Figure 6a). Subsequently, in the absence of EF-G, the L1 stalk goes through successive closed→open and open→closed transitions as the PRE complex undergoes multiple GS2→GS1 and GS1→GS2 transitions that establish a GS1⇄GS2 equilibrium (i.e., the 0.55⇄0.31 *E*_FRET_ transitions denoted by the rates *k*_GS1→GS2_ and *k*_GS2→GS1_ and the equilibrium constant *K*_eq_ = (*k*_GS1→GS2_)/(*k*_GS2→GS1_) in Figure 6d, corresponding to the GS1⇄GS2 transitions in Figure 6a). In the presence of EF-G, however, a single closed→open L1 stalk transition reports on conformational changes of the PRE complex as it undergoes EF-G binding and completes translocation (i.e., the 0.31→0.55 *E*_FRET_ transition denoted by the rate *k*_GS2→POST_ in Figures 6d and e, corresponding to the multi-step GS2→POST transition that takes place in the presence of EF-G and bridges across Figures 6a and b). Using this approach, we have successfully monitored the conformational dynamics of ribosomal complexes during individual elongation cycles^36,38–41^, including in a study of —1 frameshifting^41^.

### *SufB2* interferes with elongation complex dynamics during late steps in translocation

We began by asking whether *SufB2* alters the dynamics of elongation complexes during the earlier steps of the elongation cycle. We stopped-flow delivered *SufB2-* or *ProL*-TC to 70S ICs and recorded pre-steady-state movies during delivery, and steady-state movies 1 min post-delivery (Figures 6a, d, and f, Supplementary Figures 3, 4a, and 4b). The results showed that *k*_70S IC→GS2_, as well as *k*_GS1→GS2_, *k*_GS2→GS1_, and *K*_eq_ at 1 min post-delivery, for *SufB2*-TC were each less than 2-fold different than the corresponding value for *ProL*-TC (Supplementary Table 4). The close correspondence of these rates indicates that *SufB2*-TC is delivered to the A site, participates in peptide-bond formation, undergoes GS2 formation, and exhibits GS1→GS2 and GS2→GS1 transitions within the GS1→GS2 equilibrium in a manner that is similar to *ProL*-TC, consistent with the results of ensemble kinetic assays (Figures 3a-c, Supplementary Table 2, and Supplementary Figure 2a) and thereby strengthening our interpretation that *SufB2* uses triplet pairing in the 0-frame at the A site during the early stages of the elongation cycle that precede EF-G binding and EF-G-catalyzed translocation. Although we could not confidently detect the presence of two sub-populations of A site-bound *SufB2* in the smFRET data that might differ in their propensity of undergoing +1 frameshifting, as suggested by the results presented in Figure 5a, it is possible that the distance between our smFRET probes and/or the time spent in one of the observed FRET states are not sensitive enough to detect the structural and/or energetic differences between these sub-populations of A site-bound *SufB2*. The development of different smFRET signals and/or the use of variants of *SufB2* and/or the CCC-C motif with different propensities of undergoing +1 frameshifting may allow future smFRET investigations to identify and characterize such sub-populations.

We then investigated whether *SufB2* alters the dynamics of elongation complexes during the later steps of the elongation cycle. We stopped-flow delivered *SufB2*-or *ProL*-TC and EF-G to 70S ICs and recorded pre-steady-state movies during delivery, and steady-state movies 1, 3, 10, and 20 min post-delivery (Figures 6b, e, and g, Supplementary Figures 4c, 4d, and 5). The results showed that *k*_70S IC→GS2_ for *SufB2* and *ProL*-TC were within error of each other (Supplementary Table 5), again suggesting that *SufB2*-TC is delivered to the A site, participates in peptide-bond formation, and undergoes GS2 formation in a manner that is similar to *ProL*-TC. Notably, the *k*_70S IC→GS2_s obtained in the presence of EF-G were within error of the ones obtained in the absence of EF-G, consistent with reports that EF-G has little to no effect on the rate with which PRE complexes undergo GS1→GS2 transitions^37,42^.

Once it transitions into GS2, however, the *SufB2* PRE complex can bind EF-G^37,42^ and we find that it becomes arrested in an EF-G-bound GS2-like conformation for up to several minutes, during which it slowly undergoes a GS2→POST transition (Figure 6g, Supplementary Figure 5). While the limited number of time points did not allow rigorous determination of *k*_GS2→POST_ for the *SufB2* PRE complex, visual inspection (Figure 6g) and quantitative analysis (Supplementary Tables 5 and 6) showed that the GS2→POST reaction was complete between 3 and 10 min post-delivery (i.e., *k*_GS2→POST_ = ~0.0017-0.0060 s^-1^). Remarkably, this range of *k*_GS2→POST_ is up to 2-3 orders of magnitude lower than *k*_GS2→POST_ measured for the *ProL* PRE complex (Supplementary Table 5). It is also up to 2-3 orders of magnitude lower than *k*_GS2→POST_ for a different PRE complex measured using a different smFRET signal under the same conditions^43^ and the rate of translocation measured using ensemble rapid kinetic approaches under similar conditions^44,45^. This observation suggests that *SufB2* adopts a conformation within the EF-G-bound PRE complex that significantly impedes conformational rearrangements of the complex that are known to take place during late steps in translocation. These rearrangements include the severing of interactions between the decoding center of the 30S subunit and the anticodon-codon duplex in the A site^22–25^; forward and reverse swiveling of the ‘head’ domain of the 30S subunit^27,28^ associated with opening and closing, respectively, of the ‘E-site gate’ of the 30S subunit^26^; reverse relative rotation of the ribosomal subunits^46,47^; and opening of the L1 stalk^35,37,48^. Collectively, these dynamics facilitate movement of the tRNA ASLs and their associated codons from the P and A sites to the E and P sites of the 30S subunit.

We next explored whether *SufB2* alters the dynamics of elongation complexes after it is translocated into the P site. We prepared PRE-like complexes carrying deacylated *SufB2* or *ProL* in the P site and a vacant A site (denoted PRE^-A^ complexes) and recorded steady-state movies for the resulting GS1⇆GS2 equilibria (Figures 6c and h, Supplementary Figure 6). The results showed that *k*_GS1→GS2_ and *k*_GS2→GS1_ for the *SufB2* PRE^-A^ complex were 45% lower and 36% higher, respectively, than for the *ProL* PRE^-A^ complex, driving a 2.5-fold shift towards GS1 in the GS1⇆GS2 equilibrium (Supplementary Table 7), suggesting that *SufB2* adopts a conformation at the P site that is different from that of *ProL*. Consistent with this interpretation, a recent structural study has shown that the conformation of P site-bound *SufA6*, a +1-frameshifting tRNA with an extra nucleotide in the anticodon loop, is significantly distorted relative to a canonical tRNA^49^.

## DISCUSSION

Here we leverage the high efficiency of recoding by *SufB2* to identify the steps of the elongation cycle during which it induces +1 frameshifting at a quadruplet codon, thus answering the key questions of where, when, and how +1 frameshifting occurs. We are not aware of any other studies of +1 frameshifting that have addressed these questions as precisely. In addition to elucidating the determinants of reading-frame maintenance and the mechanisms of *SufB2*-induced +1 frameshifting, our findings reveal new principles that can be used to engineer genome recoding with higher efficiencies.

Integrating our results with the available structural, biophysical, and biochemical data on the mechanism of translation elongation results in the structure-based model for *SufB2*-induced +1 frameshifting that we present in Figure 7. In this model, POST complexes to which *SufB2* or *ProL* are delivered exhibit virtually indistinguishable conformational dynamics in the early steps of the elongation cycle, up to and including the initial GS1→GS2 transition. However, POST complexes to which *SufB2* is delivered exhibit a *k*_GS2→POST_ that is more than an order-of-magnitude slower than those to which *ProL* is delivered. Notably, *K*_GS2→POST_ comprises a series of conformational rearrangements of the EF-G-bound PRE complex that facilitate translocation of the tRNA ASLs and associated codons within the 30S subunit. These rearrangements encompass the severing of decoding center interactions with the anticodon-codon duplex in the A site^22–25^; forward and reverse head swiveling^27,28,50^ and associated opening and closing, respectively, of the E-site gate^26^; reverse relative rotation of the subunits^46,47^; and opening of the L1 stalk^35,37,48^ (steps PRE-G2 to PRE-G4, denoted with red arrows, in Figure 7). Given the importance of these rearrangements in translocation of the tRNA ASLs and their associated codons within the 30S subunit, we propose that *SufB2*-mediated perturbation of these rearrangements underlies +1 frameshifting. More specifically, because *SufB2* does not seem to impede the reverse relative rotation of the subunits or opening of the L1 stalk during the GS2→GS1 transitions within the GS1→GS2 equilibrium in the absence of EF-G (compare k_GS2→GS1_ for *SufB2*-TC vs. *Pro*L-TC in Supplementary Table 4), it most likely interferes with the severing of decoding center interactions with the anticodon-codon duplex in the A site and/or forward and/or reverse head swiveling and associated opening and/or closing, respectively, of the E-site gate. The latter rearrangement is particularly important for movement of the tRNA ASLs and their associated codons within the 30S subunit^26–28,50^, suggesting that *SufB2*-mediated perturbation of head swiveling may make the most important contribution to +1 frameshifting. Consistent with this, a recent structural study showed that upon forward head swiveling, the ASLs of the P- and A-site tRNAs can disengage from their associated codons and occupy positions similar to a partial +1 frameshift, even in the presence of a non-frameshift suppressor tRNA in the A site and the absence of EF-G^51^.

**Figure 7.**
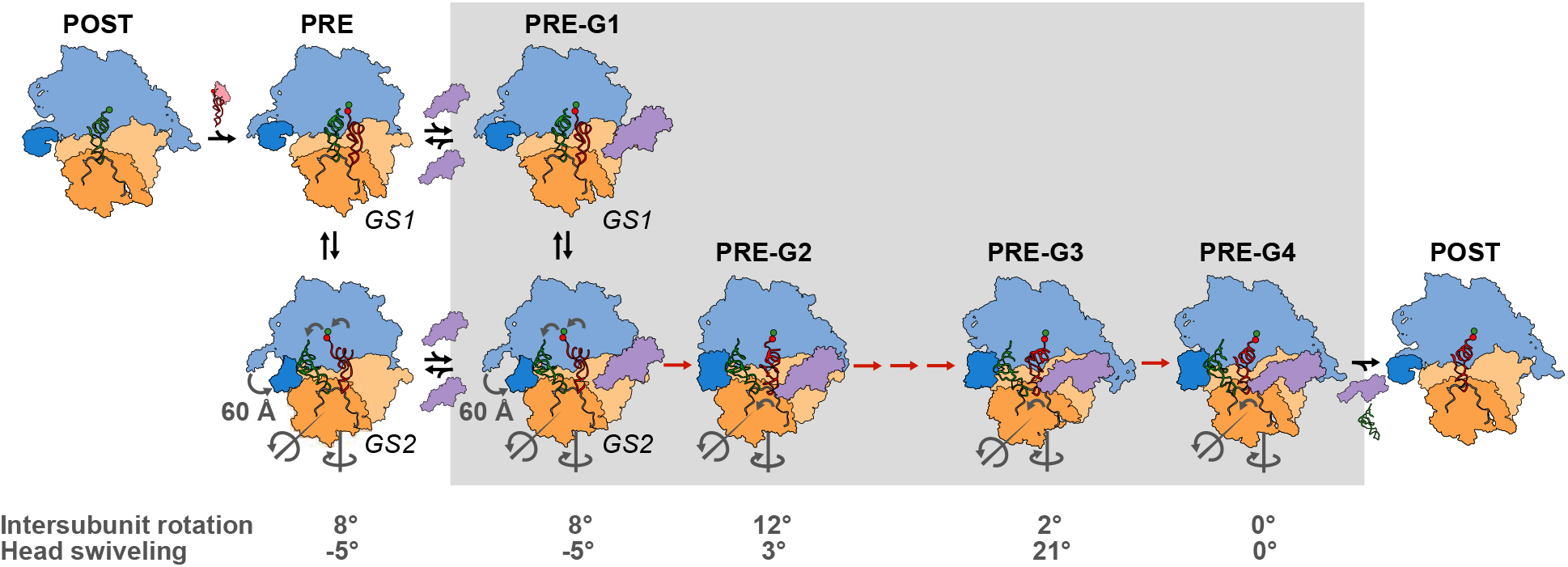
Structure-based mechanistic model for *SufB2*-induced +1 frameshifting. A *SufB2-TC* uses triplet anticodon-codon pairing in the 0-frame at a CCC-C motif, undergoes peptide-bond formation, and enables the resulting PRE complex to undergo a GS1→GS2 transition, all with rates similar to those of *ProL*-TC. During the GS1 →GS2 transition, the 30S subunit rotates relative to the 50S subunit by 8° in the counter-clockwise (+) direction along the black curved arrow; the 30S subunit head swivels relative to the 30S subunit body by 5° in the clockwise (-) direction against the black curved arrow; the L1 stalk closes by ~60 Å; and the tRNAs are reconfigured from their P/P and A/A to their P/E and A/P configurations. EF-G then binds to the PRE complex to form PRE-G1 and subsequently catalyzes a series of conformational rearrangements of the complex (PRE-G1 to PRE-G4) that encompass further counter-clockwise and clockwise rotations of the subunits; severing of decoding center interactions with the anticodon-codon duplex in the A site; counter-clockwise and clockwise swiveling of the head and the associated opening and closing of the E-site gate; opening of the L1 stalk; and reconfigurations of the tRNAs as they move from the P and A sites to the E and P sites. It is during these steps, shown in red arrows within the gray shaded box, that *SufB2* impedes forward and/or reverse swiveling of the head and the associated opening and/or closing of the E-site gate, facilitating +1 frameshifting. Next, EF-G and the deacylated tRNA dissociate from PRE-G4, leaving a POST complex ready to enter the next elongation cycle. The cartoons depicting PRE-G1(GS1) and PRE-G1(GS2) were generated using Biological Assemblies 2 and 1, respectively, of PDB entry 4V9D. Due to the lack of an A-site tRNA or EF-G in 4V9D, cartoons of the A- and P-site tRNAs from previous structures^1^ were positioned into the two assemblies using the P-site tRNAs in 4V9D as guides and a cartoon of EF-G generated from 4V7D was manually positioned near the factor binding site of the ribosomes. The cartoons depicting PRE-G2, PRE-G3, and PRE-G4 were generated from 4V7D, 4W29, and 4V5F, respectively, and colored as in Figure 6, with the head domain shown in orange.

While previous structural studies have demonstrated that +1 frameshifting tRNAs bind to the A site in the 0-frame^16,17,49^ and to the P site in the +1-frame^19^, these studies lacked EF-G and the observed structures were obtained by directly binding a deacylated +1 frameshifting tRNA to the P site. Specifically, a +1 frameshifting peptidyl-tRNA was not translocated from the A to P sites, as would be the case during an authentic translocation event. In contrast, our elucidation of the +1-frameshifting mechanism was executed in the presence of EF-G and is based on extensive comparison of the kinetics with which *SufB2* and *ProL* undergo individual reactions of the elongation cycle (i.e., aa-tRNA selection, peptide-bond formation, and translocation) and the associated conformational rearrangements of the elongation complex. Additionally, all of our in vitro biochemical assays, and most of our ensemble rapid kinetics assays were performed under the conditions in which the A site is always occupied by an aa-or peptidyl-tRNA, leaving no chance of a vacant A site. Therefore, the +1 frameshifting mechanism we present here is distinct from that presented by Farabaugh and co-workers^13^, in which the ribosome is stalled due to a vacant A site, thus giving the +1-frameshifting-inducing tRNA at the P site an opportunity to rearrange into the +1-frame. The fact that all well-characterized +1-frameshifting tRNAs contain an extra nucleotide in the anticodon loop, despite differences in their primary sequences, the amino acids they carry, and whether the extra nucleotide is inserted at the 3’- or 5’-sides of the anticodon, suggests that the results we report here for *SufB2* are likely applicable to other +1-frameshifting tRNAs with an expanded anticodon loop.

While an expanded anticodon loop is a strong feature associated with +1 frameshifting, it is not associated with −1 frameshifting, which instead is typically induced by structural barriers in the mRNA that stall a translating ribosome from moving forward, thus providing the ribosome with an opportunity to shift backwards in the −1 direction^10,52^. Given the unique role of the expanded anticodon loop in +1 frameshifting, here we have identified the determinants that drive the ribosome to shift in the +1 direction. We show that *SufB2* exclusively uses the triplet-slippage mechanism of +1 frameshifting in the m^1^G37+ condition, but that it explores other mechanisms (e.g., quadruplet-pairing) in the m^1^G37-condition during translocation from the A site to the P site. Under conditions that only permit the triplet-slippage mechanism (e.g., in the presence of m^1^G37), *SufB2* exhibits a relatively low +1-frameshifting efficiency of ~30%, whereas under conditions that permit quadruplet-pairing during translocation (e.g., in the absence of m^1^G37), it exhibits a relatively high +1-frameshifting efficiency of ~90% (Figures 4c-f, 5a). This feature is observed in various sequence contexts. One advantage of a quadruplet-pairing mechanism during translocation is that it would enhance the thermodynamic stability of anticodon-codon pairing during the large EF-G-catalyzed conformational rearrangements that PRE complexes undergo during translocation to form POST complexes. Nonetheless, *SufB2* is naturally methylated with m^1^G37 (Figure 1c), indicating that it makes exclusive use of the triplet-slippage mechanism in vivo. This mechanism is likely also exclusively used in vivo by all other +1-frameshifting tRNAs that have evolved from canonical tRNAs to retain a purine at position 37, which is almost universally post-transcriptionally modified to block quadruplet-pairing mechanisms.

The key insight from this work suggests an entirely novel pathway to increase the efficiency of genome recoding at quadruplet codons. While initial success in genome recoding has been achieved by engineering the anticodon-codon interactions of a +1-frameshifting-inducing tRNA at the A site^6,53^, or by engineering a new bacterial genome with a minimal set of codons for all amino acids^54^, we suggest that efforts to engineer the ‘neck’ structural element of the 30S subunit that regulates head swiveling would be as, or even more, effective. This can be achieved by screening for 30S subunit variants that exhibit high +1-frameshifting efficiencies mediated by +1-frameshifting tRNAs at quadruplet codons while preserving 0-frame translation by canonical tRNAs at triplet codons. Specifically, head swiveling is driven by the synergistic action of two hinges within the 16S ribosomal RNA elements that comprise the 30S subunit neck^55^. Hinge 1 is composed of two G-U wobble base pairs that are separated by a bulged G within helix 28 (h28), while hinge 2 is composed of a GACU linker between h34 and h35/36 within a three-helical junction with h38. Co-engineering these two hinges by directed evolution should identify such 30S subunit variants. To complement the directed evolution approach, we suggest that our recently developed time-resolved cryogenic electron microscopy (TR cryo-EM) method^56,57^ can be used to obtain structures of *SufB2* and *ProL* in EF-G-bound PRE complexes captured in intermediate states of translocation. Such cryo-EM structures would help further define how the two hinges that control head swiveling are differentially modulated during translocation of *SufB2* vs. *ProL* to provide a structure-based roadmap for engineering them. In addition, detailed comparison of such structures would offer the opportunity to identify ribosomal structural elements beyond the two hinges that play a role in +1 frameshifting and can thus serve as additional targets for engineering. Furthermore, antibiotics that bind to the 30S subunit and act as translocation modulators can be exploited to further increase the +1-frameshifting efficiency at a quadruplet codon with either wildtype or highly efficient 30S subunit variants. Implemented in combination and integrated into a recently described in vivo ‘designer organelle’ strategy^58^, these approaches should provide a novel and powerful platform for increasing the efficiency of genome recoding at quadruplet codons with minimal off-target effects.

## METHODS

### Construction of *E. coli* strains

*E. coli* strains that expressed a plasmid-borne *ProL* or *SufB2* for isolation of native-state tRNAs were made in a *ProL-KO* strain, which was constructed by inserting the Kan-resistance (Kan-R) gene, amplified by PCR primers from pKD4, into the *ProL* locus of *E. coli* BL21(DE3) using the l-Red recombination method^59^, followed by removal of the Kan-R gene using FLP recombination^30^. The pKK223-*SufB2* plasmid was made by site-directed mutagenesis to introduce G37a into the pKK223-*ProL* plasmid^29^. *E. coli* strains that expressed *ProL* or *SufB2* from the chromosome as an isogenic pair for reporter assays were made using the l-Red technique^30^. To construct the *E. coli SufB2* strain, the *SufB2* gene was PCR-amplified from pKK223-*SufB2*, and the 5’ end of the amplified gene was joined with Kan-R (from pKD4) by PCR using reverse-2 primer, while the 3’ end was homologous to the *ProL* 3’ flanking region. The PCR-amplified *SufB2-Kan* product was used to replace *ProL* in l-Red expressing cells. An isogenic counterpart strain expressing *ProL-Kan* was also made. These *ProL-Kan* and *SufB2-Kan* loci were independently transferred to the *trmD-KO* strain^29^ by P1 transduction, followed by pCP20-dependent FLP recombination, generating the isogenic pair of *ProL* and *SufB2* strains in the *trmD-KO* background. These strains were transformed with *pKK223-3-lacZ* reporter plasmid that has the CCC-C motif at the 2^nd^ codon position of the *lacZ* gene, and the β-Gal activity was measured^29^. All primer sequences used in this work are shown in Supplementary Table 1.

### Preparation of translation components for ensemble biochemical experiments

The mRNA used for most in vitro translation reactions is shown below, including the Shine-Dalgarno sequence, the AUG start codon, and the CCC-C motif:

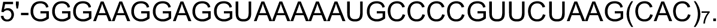

Variants of this mRNA had a base substitution in the CCC-C motif. All mRNAs were transcribed from double-stranded DNA templates with T7 RNA polymerase and purified by gel electrophoresis. *E. coli* strains over-expressing native-state tRNA^fMet^, tRNA^Arg^ (anticodon ICG, where I = inosine), and tRNA^Val^ (anticodon U*AC, where U* = cmo^5^U) were grown to saturation and were used to isolate total tRNA. The over-expressed tRNA species in each total tRNA sample was aminoacylated by the cognate aminoacyl-tRNA synthetase and used directly in the TC formation reaction and subsequent TC delivery to 70S ICs or POST complexes. *E. coli* tRNA^ser^ (anticodon ACU) was prepared by in vitro transcription. Aminoacyl-tRNAs with the cognate proteinogenic amino acid were prepared using the respective aminoacyl-tRNA synthetase and those with a non-proteinogenic amino acid were prepared using the dFx Flexizyme and the 3,5-dinitobenzyl ester (DBE) of the respective amino acid (Supplementary Figure 1). Aminoacylation and formylation of tRNA^fMet^ were performed in a one-step reaction in which formyl transferase and the methyl donor 10-formyltetrahydrofolate were added to the aminoacylation reaction^29^. Aminoacyl-tRNAs were stored in 25 mM sodium acetate (NaOAc) (pH 5) at −70 °C, as were six-His-tagged *E. coli* initiation and elongation factors and tight-coupled 70S ribosomes isolated from *E. coli* MRE600 cells. Recombinant His-tagged *E. coli* EF-P bearing a β-lysyl-K34 was expressed and purified from cells co-expressing *efp, yjeA*, and *yjeK* and stored at −20 °C^29^.

### Preparation of translation components for smFRET experiments

30S subunits and 50S subunits lacking ribosomal proteins bL9 and uL1 were purified from a previously described bL9-uL1 double deletion *E. coli* strain^35,60^ using previously described protocols^35,37,60^. A previously described single-cysteine variant of bL9 carrying a Gln-to-Cys substitution mutation at residue position 18 (bL9(Q18C))^35^ and a previously described single-cysteine variant of uL1 carrying a Thr-to-Cys substitution mutation at residue position 202 (uL1(T202C))^35,37^ were purified, labeled with Cy3- and Cy5-maleimide, respectively, to generate bL9(Cy3) and uL1(Cy5), and reconstituted into the 50S subunits lacking bL9 and uL1 following previously described protocols^35^. The reconstituted bL9(Cy3)- and uL1(Cy5)-labeled 50S subunits were then re-purified using sucrose density gradient ultracentrifugation^35,43^. 50S subunits lacking bL9(Cy3) and/or uL1(Cy5) or harboring unlabeled bL9 and/or uL1 do not generate bL9(Cy3)-uL1(Cy5) smFRET signals and therefore do not affect data collection or analysis. Previously, we have shown that 70S ICs formed with these bL9(Cy3)- and uL1(Cy5)-containing 50S subunits can undergo peptide-bond formation and two rounds of translocation elongation with similar efficiency as 70S ICs formed with wild-type 50S subunits^35^.

The sequence of the mRNA used for assembling ribosomal complexes for smFRET studies is shown below, including the Shine-Dalgarno sequence, the AUG start codon, and the CCC-C motif:

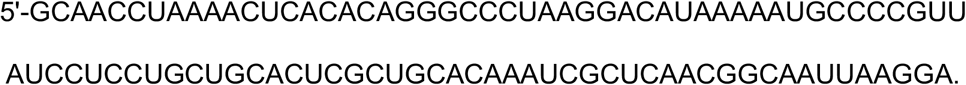

The mRNA was synthesized by in vitro transcription using T7 RNA polymerase, and then hybridized to a previously described 3’-biotinylated DNA oligonucleotide (Supplementary Table 1) that was complementary to the 5’ end of the mRNA and was chemically synthesized by Integrated DNA Technologies^60^. Hybridized mRNA:DNA-biotin complexes were stored in 10 mM Tris-OAc (pH = 7.5 at 37 °C), 1 mM EDTA, and 10 mM KCl at −80 °C until they were used in ribosomal complex assembly. Aminoacylation and formylation of tRNA^fMet^ (purchased from MP Biomedicals) was achieved simultaneously using *E. coli* methionyl-tRNA synthetase and *E. coli* formylmethionyl-tRNA formyltransferase^60^. Expression and purification of IF1, IF2, IF3, EF-Tu, EF-Ts, and EF-G were following previously published procedures^60^.

### Preparation and purification of *SufB2* and *ProL*

Native-state *SufB2* was isolated from a derivative of *E. coli* JM109 lacking the endogenous *ProL*, but expressing *SufB2* from the pKK223-3 plasmid (Supplementary Table 1), while native-state *ProL* was purified from total tRNA isolated from *E. coli* JM109 cells over-expressing *ProL* from the pKK223-3 plasmid. The *ProL-KO* strain lacking the endogenous *ProL* was described previously^30^. Each native-state tRNA was isolated by a biotinylated capture probe attached to streptavidin-derivatized Sepharose beads^29^. G37-state *SufB2* and *ProL* were also prepared by in vitro transcription. Each primary transcript contained a ribozyme domain on the 5’-side of the tRNA sequence, which self-cleaved to release the tRNA. m^1^G37-state *SufB2* and *ProL* were prepared by TrmD-catalyzed and *S*-adenosyl methionine (AdoMet)-dependent methylation of each G37-state tRNA. Due to the lability of the aminoacyl linkage to Pro, stocks of *SufB2* and *ProL* aminoacylated with Pro were either used immediately or stored no longer than 2-3 weeks at −70 °C in 25 mM NaOAc (pH 5.0).

### Primer extension inhibition assays

Primer extension inhibition analyses of native-, G37-, and m^1^G37-state *SufB2* and *ProL* were performed as described^30^. A DNA primer complementary to the sequence of C41 to A57 of *SufB2* and *ProL* was chemically synthesized, ^32^P-labeled at the 5’-end by T4 polynucleotide kinase, annealed to each tRNA, and was extended by Superscript III reverse transcriptase (Invitrogen) at 200 units/μL with 6 μM each dNTP in 50 mM Tris-HCl (pH 8.3), 3 mM MgCl_2_, 75 mM KCl, and 1 mM DTT at 55 °C for 30 min, and terminated by heating at 70 °C for 15 min. Extension was quenched with 10 mM EDTA and products of extension were separated by 12% denaturing polyacrylamide gel electrophoresis (PAGE/7M urea) and analyzed by phosphorimaging. In these assays, the length of the read-through cDNA is 54-55 nucleotides, as in the case of the G37-state *SufB2* and *ProL*, whereas the length of the primer-extension inhibited cDNA products is 21-22 nucleotides, as in the case of the m^1^G37-state and native-state.

### RNase T1 cleavage inhibition assays

RNase T1 cleaves on the 3’-side of G, but not m^1^G. Cleavage of tRNAs was performed as previously described^29^. Each tRNA (1 μg) was 3’-end labeled using *Bacillus stearothermophilus* CCA-adding enzyme (10 nM) with [α-^32^P]ATP at 60 °C in 100 mM glycine (pH 9.0) and 10 mM MgCl_2_. The labeled tRNA was digested by RNase T1 (Roche, cat # 109193) at a final concentration of 0.02 units/μL for 20 min at 50 °C in 20 mM sodium citrate (pH 5.5) and 1 mM ethylene diamine tetraacetic acid (EDTA). The RNA fragments generated from cleavage were separated by 12% PAGE/7M urea along with an RNA ladder generated by alkali hydrolysis of the tRNA of interest. Cleavage was analyzed by phosphorimaging.

### Methylation assays

Pre-steady-state assays under single-turnover conditions^61^ were performed on a rapid quench-flow apparatus (Kintek RQF-3). The tRNA substrate was heated to 85 °C for 2.5 min followed by addition of 10 mM MgCl_2_, and slowly cooled to 37 °C in 15 min. *N*^1^-methylation of G37 in the pre-annealed tRNA (final concentration 1 μM) was initiated with the addition of *E. coli* TrmD (10 μM) and [^3^H]-AdoMet (Perkin Elmer, 4200 DPM/pmol) at a final concentration of 15 μM in a buffer containing 100 mM Tris-HCl (pH 8.0), 24 mM NH_4_Cl, 6 mM MgCl_2_, 4 mM DTT, 0.1 mM EDTA, and 0.024 mg/mL BSA in a reaction of 30 μL. The buffer used was optimized for TrmD in order to evaluate its in vitro activity^61^. Reaction aliquots of 5 μL were removed at various time points and precipitated in 5% (w/v) trichloroacetic acid (TCA) on filter pads for 10 min twice. Filter pads were washed with 95% ethanol twice, with ether once, air dried, and measured for radioactivity in an LS6000 scintillation counter (Beckman). Counts were converted to pmoles using the specific activity of the [^3^H]-AdoMet after correcting for the signal quenching by filter pads. In these assays, a negative control was always included, in which no enzyme was added to the reaction^61^, and signal from the negative control was subtracted from signal of each sample for determining the level of methylation.

### Aminoacylation assays

Each *SufB2* or *ProL* tRNA was aminoacylated with Pro by a recombinant *E. coli* ProRS expressed from the plasmid pET22 and purified from *E. coli* BL21 (DE3)^62^. Each tRNA was heat-denatured at 80 °C for 3 min, and re-annealed at 37 °C for 15 min. Aminoacylation under pre-steady state conditions was performed at 37 °C with 10 μM tRNA, 1 μM ProRS, and 15 μM [^3^H]-Pro (Perkin Elmer, 7.5 Ci/mmol) in a buffer containing 20 mM KCl, 10 mM MgCl_2_, 4 mM dithiothreitol (DTT), 0.2 mg/mL bovine serum albumin (BSA), 2 mM ATP (pH 8.0), and 50 mM Tris-HCl (pH 7.5) in a reaction of 30 μL. Reaction aliquots of 5 μL were removed at different time intervals and precipitated with 5% (w/v) TCA on filter pads for 10 min twice. Filter pads were washed with 95% ethanol twice, with ether once, air dried, and measured for radioactivity in an LS6000 scintillation counter (Beckman). Counts were converted to pmoles using the specific activity of the [^3^H]-Pro after correcting for signal quenching by filter pads.

### Cell-based +1-frameshifting reporter assays

Isogenic *E. coli* strains expressing chromosomal copies of *SufB2* or *ProL* were created in a previously developed *trmD-knockdown (trmD-KD)* background, in which the chromosomal *trmD* is deleted but cell viability is maintained through the arabinose-induced expression of a plasmid-borne *trm5*, the human counterpart of *trmD*^29,30^ that is competent for m^1^G37 synthesis to support bacterial growth (Supplementary Table 1). Due to the essentiality of *trmD* for cell growth, a simple knock-out cannot be made. We chose human Trm5 as the maintenance protein in the *trmD-KD* background, because this enzyme is rapidly degraded in *E. coli* once its expression is turned off to allow immediate arrest of m^1^G37 synthesis. In the isogenic *SufB2* and *ProL* strains, the level of m^1^G37 is determined by the concentration of the added arabinose in a cellular context that expresses *ProM* as the only competing tRNA^pro^ species. In the m^1^G37+ condition, where arabinose was added to 0.2% in the medium, tRNA substrates of *N*^1^-methylation were confirmed to be 100% methylated by mass spectrometry, whereas in the m^1^G37-condition, where arabinose was not added to the medium, tRNA substrates of *N*^1^-methylation were confirmed to be 20% methylated by mass spectrometry^30^. Each strain was transformed with the pKK223-3 plasmid expressing an mRNA with a CCC-C motif at the 2^nd^ codon position of the reporter *lacZ* gene. To simplify the interpretation, the natural AUG codon at the 5^th^ position of *lacZ* was removed. A +1 frameshift at the CCC-C motif would enable expression of *lacZ*. The activity of β-Gal was directly measured from lysates of cells grown in the presence or absence of 0.2% arabinose to induce or not induce, respectively, the plasmid-borne human *trm5*. In these assays, decoding of the CCC-C codon motif would be mediated by *SufB2* and *ProM* in the *SufB2* strain, and would be mediated by *ProL* and *ProM* in the *ProL* strain. Due to the presence of *ProM* in both strains, there would be no vacancy at the CCC-C codon motif.

### Cell-based +1 frameshifting *lolB* assays

To quantify the +1-frameshifting efficiency at the CCC-C motif at the 2^nd^ codon position of the natural *lolB* gene, the ratio of protein synthesis of *lolB* to *cysS* was measured by Western blots. Overnight cultures of the isogenic strains expressing *SufB2* or *ProL* were separately inoculated into fresh LB media in the presence or absence of 0.2% arabinose and were grown for 4 h to produce the m^1^G37+ and m^1^G37-conditions, respectively. Cultures were diluted 10-to 16-fold into fresh media to an optical density (OD) of ~0.1 and grown for another 3 h. Cells were harvested and 15 μg of total protein from cell lysates was separated on 12% SDS-PAGE and probed with rabbit polyclonal primary antibodies against LolB (at a 10,000 dilution) and against CysRS (at a 20,000 dilution), followed by goat polyclonal anti-rabbit IgG secondary antibody (Sigma-Aldrich, #A0545). The ratio of protein synthesis of *lolB* to *cysS* was quantified using Super Signal West Pico Chemiluminescent substrate (Thermo Fischer) in a Chemi-Doc XR imager (Bio-Rad) and analyzed by Image Lab software (Bio-Rad, SOFT-LIT-170-9690-ILSPC-V-6-1). To measure the +1-frameshifting efficiency, we measured the ratio of protein synthesis of *lolB* to *cysS* for each tRNA in each condition, and we normalized the observed ratio in the control sample (i.e., *ProL* in the m^1^G37+ condition) to 1.0, indicating that protein synthesis of these two genes was in the 0-frame and no +1 frameshifting. A decrease of this ratio was interpreted as a proxy of +1 frameshifting at the CCC-C motif at the 2^nd^ codon position of *lolB*. From the observed ratio of each sample in each condition, we calculated the +1 frameshifting efficiency relative to the control sample.

### Cell-free PURExpress in vitro translation assays

The *folA* gene, provided as part of the *E. coli* PURExpress (New England BioLabs) in vitro translation system, was modified by site-directed mutagenesis to introduce a CCC-C motif into the 5^th^ codon position. If *SufB2* induced +1 frameshifting at this motif, a full-length DHFR would be made, whereas if *SufB2* failed to do so, a C-terminal truncated fragment (ΔC) would be made due to premature termination of protein synthesis. Because *SufB2* has no orthogonal tRNA synthetase for aminoacylation with a non-proteinogenic amino acid, we used the Flexizyme ribozyme technology^32^ for this purpose. Coupled in vitro transcription-translation of the modified *E. coli folA* gene containing the CCC-C motif at the 5^th^ codon position was conducted in the presence of [^35^S]-Met using the PURExpress system. SDS-PAGE analysis was used to detect [^35^S]-Met-labeled polypeptides, which included the full-length DHFR, the ΔC fragment, and a ΔN fragment that likely resulted from initiation of translation at a cryptic site downstream from the CCC-C motif (Figure 2d). The fraction of the fulllength *folA* gene product, the ΔC fragment, and the ΔN fragment was calculated from the amount of each in the sum of all three products. We attribute the overall low recoding efficiency (0.5 - 5.0%) as arising from a combination of the rapid hydrolysis of the prolyl linkage, which is the least stable among aminoacyl linkages^63^, and the lack of *SufB2* re-acylation in the PURExpress system. In these assays, each tRNA was tested in the G37-state and each was normalized by the flexizyme aminoacylation efficiency, which was ~30% for Pro and Pro analogues. The PURExpress contained all natural *E. coli* tRNAs, such that the CCC-C codon motif would not have a chance of vacancy even when a specific CCC-reading tRNA was absent.

### Rapid kinetic GTPase assays

Ensemble GTPase assays were performed using the codon-walk approach, in which an *E. coli* in vitro translation system composed of purified components is supplemented with the requisite tRNAs and translation factors to interrogate individual steps of the elongation cycle. Programmed with a previously validated synthetic AUG-CCC-CGU-U mRNA template^29,34^, a 70S IC was assembled that positioned the AUG start codon and an initiator fMet-tRNA^fMet^ at the P site and the CCC-C motif at the A site. Reactions to monitor the EF-Tu-dependent hydrolysis of GTP during delivery and accommodation of a TC to the A site were conducted at 20 °C in a buffer containing 50 mM Tris-HCl (pH 7.5), 70 mM NH4Cl, 30 mM KCl, 7 mM MgCl_2_, 1 mM DTT, and 0.5 mM spermidine^29^. Each TC was formed by incubating EF-Tu with 8 nM [γ-^32^P]-GTP (6000 C_i_/mmole) for 15 min at 37 °C, after which aminoacylated *SufB2* or *ProL* was added and the incubation continued for 15 min at 4 °C. Unbound [γ-^32^P]-GTP was removed from the TC solution by gel filtration through a spin cartridge (CentriSpin-20; Princeton Separations). Equal volumes of each purified TC and a solution of 70S ICs were rapidly mixed in the RQF-3 Kintek chemical quench apparatus^29^. Final concentrations in these reactions were 0.5 μM for the 70S IC; 0.8 μM for mRNA; 0.65 μM each for IFs 1,2, and 3; 0.65 μM for fMet-tRNA^Met^; 1.8 μM for EF-Tu; 0.4 μM for aminoacylated *SufB2* or *ProL*; and 0.5 mM for cold GTP. The yield of GTP hydrolysis and *k*_GTP,obs_ upon rapid mixing of each TC with excess 70S ICs were measured by removing aliquots of the reaction at defined time points, quenching the aliquots with 40% formic acid, separating [γ-^32^P] from [γ-^32^P]-GTP using thin layer chromatography (TLC), and quantifying the amount of each as a function of time using phosphorimaging^29^. We adjusted reaction conditions such that the *k*_GTP,obs_ increased linearly as a function of 70S IC concentration.

### Rapid kinetic di- and tripeptide formation assays

Di- and tripeptide formation assays were performed using the codon-walk approach described above in 50 mM Tris-HCl (pH 7.5), 70 mM NH_4_Cl, 30 mM KCl, 3.5 mM MgCl_2_, 1 mM DTT, 0.5 mM spermidine, at 20 °C unless otherwise indicated^29^. 70S ICs were formed by incubating 70S ribosomes, mRNA, [^35^S]-fMet-tRNA^fMet^, and IFs 1,2, and 3, and GTP, for 25 min at 37 °C in the reaction buffer. Separately, TCs were formed in the reaction buffer by incubating EF-Tu and GTP for 15 min at 37 °C followed by adding the requisite aa-tRNAs and incubating in an ice bath for 15 min. In dipeptide formation assays, 70S ICs templated with the specified variants of an AUG-NNN-NGU-U mRNA were mixed with *SufB2-TC* or *ProL*-TC. fMP formation was monitored in an RQF-3 Kintek chemical quench apparatus. In tripeptide formation assays, 70S ICs templated with the specified variants of the AUG-NCC-NGU-U mRNA were mixed, either in one step or in two steps, with equimolar mixtures of *SufB2*-, tRNA^Val^ (anticodon U*AC, where U* = cmo^5^U)-, and tRNA^Arg^ (anticodon ICG, where I = inosine)-TCs and EF-G. Formation of fMPV and fMPR were monitored in an RQF-3 Kintek chemical quench apparatus. Tripeptide formation assays with one-step delivery of TCs were initiated by rapidly mixing the 70S IC with two or more of the TCs in the RQF-3 Kintek chemical quench apparatus. Final concentrations in these reactions were 0.37 μM for the 70S IC; 0.5 μM for mRNA; 0.5 μM each for IFs 1,2, and 3; 0.25 μM for [^35^S]-fMet-tRNA^fMet^; 2.0 μM for EF-G; 0.75 μM for EF-Tu for each aa-tRNA; 0.5 μM each for the aa-tRNAs; and 1 mM for GTP. For tripeptide formation assays with one-step delivery of G37-state *SufB2-*, tRNA^Val^-, and tRNA^Arg^-TCs to the 70S ICs, the yield of fMPV and *k*_fMPV,obs_ report on the activity of ribosomes that shifted to the +1-frame, whereas the yield of fMPR and *k*_fMPR,obs_ report on the activity of ribosomes that remained in the 0-frame^29,34^.

We chose G37-state *SufB2* to maximize its +1-frameshifting efficiency but native-state tRNA^Val^ and tRNA^Arg^ to prevent them from undergoing unwanted frameshifting (note that, for simplicity, we have not denoted the aminoacyl or dipeptidyl moieties of the tRNAs). Tripeptide formation assays with two-step delivery of TCs^29^ were performed in a manner similar to those with one-step delivery of TCs, except that the 70S ICs were incubated with a *SufB2-* or *ProL*-TC and 2.0 μM EF-G for 0.5-10 min, as specified, followed by manual addition of an equimolar mixture of tRNA^Arg^- and tRNA^Val^-TCs. Reactions were conducted at 20 °C unless otherwise specified, and were quenched by adding concentrated KOH to 0.5 M. After a brief incubation at 37 °C, aliquots of 0.65 μL were spotted onto a cellulose-backed plastic TLC sheet and electrophoresed at 1000 V in PYRAC buffer (62 mM pyridine, 3.48 M acetic acid, pH 2.7) until the marker dye bromophenol blue reached the water-oil interface at the anode^29^. The position of the origin was adjusted to maximize separation of the expected oligopeptide products. The separation of unreacted [^35^S]-fMet and each of the [^35^S]-fMet-peptide products was visualized by phosphorimaging and quantified using ImageQuant (GE Healthcare) and kinetic plots were fitted using Kaleidagraph (Synergy software).

### Assembly and purification of 70S ICs, TCs, POST, and PRE^-A^ complexes for use in smFRET experiments

70S ICs were assembled in a manner analogous to those for the ensemble rapid kinetic studies described above, except that the mRNA containing an AUG-CCC-CGU-U coding sequence was 5’-biotinylated and the 50S subunits were labeled with bL9(Cy3) and uL1(Cy5). More specifically, 70S ICs were assembled in three steps. First, 15 pmol of 30S subunits, 27 pmol of IF1, 27 pmol of IF2, 27 pmol of IF3, 18 nmol of GTP, and 25 pmol of biotin-mRNA in 7 μL of Tris-Polymix Buffer (50 mM Tris-(hydroxymethyl)-aminomethane acetate (Tris-OAc) (pH_25°C_ = 7.0), 100 mM KCl, 5 mM NH_4_OAc, 0.5 mM Ca(OAc)_2_, 0.1 mM EDTA, 10 mM 2-mercaptoethanol (BME), 5 mM putrescine dihydrochloride, and 1 mM spermidine (free base)) at 5 mM Mg(OAc)_2_ were incubated for 10 min at 37 °C. Then 20 pmol of fMet-tRNA^fMet^ in 2 μL of 10 mM KOAc (pH = 5) was added to the reaction, followed by an additional incubation of 10 min at 37 °C. Finally, 10 pmol of bL9(Cy3)- and uL1(Cy5)-labeled 50S subunits in 1 μL of Reconstitution Buffer (20 mM Tris-HCl (pH_25°C_ = 7.8), 8 mM Mg(OAc)_2_, 150 mM NH4Cl, 0.2 mM EDTA, and 5 mM BME) was added to the reaction to give a final volume of 10 μL, followed by a final incubation of 10 min at 37 °C. The reaction was then adjusted to 100 μL with Tris-Polymix Buffer at 20 mM Mg(OAc)_2_, loaded onto a 10-40% (w/v) sucrose gradient prepared in Tris-Polymix Buffer at 20 mM Mg(OAc)_2_, and purified by sucrose density gradient ultracentrifugation to remove any free mRNA, IFs, and fMet-tRNA^fMet^. Purified 70S ICs were aliquoted, flash frozen in liquid nitrogen, and stored at −80 °C until use in smFRET experiments.

TCs were prepared in two steps. First, 300 pmol of EF-Tu and 200 pmol of EF-Ts in 8 μL of Tris-Polymix Buffer at 5 mM Mg(OAc)_2_ supplemented with GTP Charging Components (1 mM GTP, 3 mM phosphoenolpyruvate, and 2 units/mL pyruvate kinase) were incubated for 1 min at 37 °C. Then, 30 pmol of aa-tRNA in 2 μL of 25 mM NaOAc (pH = 5) was added to the reaction, followed by an additional incubation of 1 min at 37 °C. This results in a TC solution with a final volume of 10 μL that was then stored on ice until used for smFRET experiments.

To prepare PRE^-A^ complexes, we first needed to assemble POST complexes. POST complexes were assembled by first preparing a 10-μL solution of 70S IC and a 10-μL solution of TC as described above. Separately, a solution of GTP-bound EF-G was prepared by incubating 120 pmol EF-G in 5 μL of Tris-Polymix Buffer at 5 mM Mg(OAc)_2_ supplemented with GTP Charging Components for 2 min at room temperature. Then 10 μL of the 70S IC, 10 μL of the TC, and 2.5 μL the GTP-bound EF-G solution were mixed, and incubated for 5 min at room temperature and for additional 5 min on ice. The resulting POST complex was diluted by adjusting the reaction volume to 100 μL with Tris-Polymix Buffer at 20 mM Mg(OAc)_2_ and purified via sucrose density gradient ultracentrifugation as described above for the 70S ICs. Purified POST complexes were aliquoted, flash frozen in liquid nitrogen, and stored at −80 °C until use in smFRET experiments. PRE^-A^ complexes were then generated by mixing 3 μL of POST complex, 2 μL of a 10 mM puromycin solution (prepared in Nanopure water and filtered using a 0.22 μm filter), and 15 μL of Tris-Polymix Buffer at 15 mM Mg(OAc)_2_ and incubating the mixture for 10 min at room temperature. PRE^-A^ complexes were used for smFRET experiments immediately upon preparation.

### smFRET imaging using total internal reflection fluorescence (TIRF) microscopy

70S ICs or PRE^-A^ complexes were tethered to the PEG/biotin-PEG-passivated and streptavidin-derivatized surface of a quartz microfluidic flowcell via a biotin-streptavidin-biotin bridge between the biotin-mRNA and the biotin-PEG^37,43^. Untethered 70S ICs or PRE^-A^ complexes were removed from the flowcell, and the flowcell was prepared for smFRET imaging experiments, by flushing it with Tris-Polymix Buffer at 15 mM Mg(OAc)_2_ supplemented with an Oxygen-Scavenging System (2.5 mM protocatechuic acid (pH = 9) (Sigma Aldrich) and 250 nM protocatechuate-3,4-dioxygenase (pH = 7.8) (Sigma Aldrich))^64^ and a Triplet-State-Quencher Cocktail (1 mM 1,3,5,7-cyclooctatetraene (Aldrich) and 1 mM 3-nitrobenzyl alcohol (Fluka))^65^.

Tethered 70S ICs or PRE^-A^ complexes were imaged at single-molecule resolution using a laboratory-built, wide-field, prism-based total internal reflection fluorescence (TIRF) microscope with a 532-nm, diode-pumped, solid-state laser (Laser Quantum) excitation source delivering a power of 16-25 mW as measured at the prism to ensure the same power density on the imaging plane. The Cy3 and Cy5 fluorescence emissions were simultaneously collected by a 1.2 numerical aperture, 60×, water-immersion objective (Nikon) and separated based on wavelength using a two-channel, simultaneous-imaging system (Dual View™, Optical Insights LLC). The Cy3 and Cy5 fluorescence intensities were recorded using a 1024 × 1024 pixel, back-illuminated electron-multiplying charge-coupled-device (EMCCD) camera (Andor iXon Ultra 888) operating with 2 × 2 pixel binning at an acquisition time of 0.1 seconds per frame controlled by software μManager 1.4. This microscope allows direct visualization of thousands of individual 70S ICs or PRE^-A^ complexes in a field-of-view of 115 × 230 μm^2^. Each movie was composed of 600 frames in order to ensure that the majority of the fluorophores in the field-of-view were photobleached within the observation period. For stopped-flow experiments using tethered 70S ICs, we delivered 0.25 μM of G37-state *SufB2-* or *ProL*-TC in the absence of EF-G or, when specified, in the presence of a 2 μM saturating concentration of EF-G. Stopped-flow experiments proceeded by recording an initial pre-steady-state movie of a field-of-view that captured conformational changes taking place during delivery followed by recording of one or more steady-state movies of different fields-of-view that captured conformational changes taking place the specified number of minutes post-delivery.

### Analysis of smFRET experiments

For each TIRF microscopy movie, we identified fluorophores, aligned Cy3 and Cy5 imaging channels, and generated fluorescence intensity vs. time trajectories for each pair of Cy3 and Cy5 fluorophores using custom-written software (manuscript in preparation; Jason Hon, Colin Kinz-Thompson, Ruben L. Gonzalez) as described previously^66^. For each time point, Cy5 fluorescence intensity values were corrected for Cy3 bleedthrough by subtracting 5% of the Cy3 fluorescence intensity value in the corresponding Cy3 fluorescence intensity vs. time trajectory. *E*_FRET_ vs. time trajectories were generated by using the Cy3 fluorescence intensity (*I*_cy3_) and the bleedthrough-corrected Cy5 fluorescence intensity (*I*_cy5_) from each aligned pair of Cy3 and Cy5 fluorophores to calculate the *E*_FRET_ value at each time point using *E*_FRET_ = (*I*_cy5_ / (*I*_cy5_ + *I*_cy3_)).

For both pre-steady-state and steady-state movies (Figures 6d–6h and Supplementary Figures 3, 5, and 6, Supplementary Tables 4-7), an *E*_FRET_ vs. time trajectory was selected for further analysis if all of the transitions in the fluorescence intensity vs. time trajectory were anti-correlated for the corresponding, aligned pair of Cy3 and Cy5 fluorophores, and the Cy3 fluorescence intensity vs. time trajectory underwent single-step Cy3 photobleaching, demonstrating it arose from a single ribosomal complex. In the case of pre-steady-state movies (Figures 6d–6g, Supplementary Figures 3 and 5 and Tables 4-6), *E*_FRET_ vs. time trajectories had to meet two additional criteria in order to be selected for further analysis: (i) *E*_FRET_ vs. time trajectories had to be stably sampling *E*_FRET_ = 0.55 prior to TC delivery, thereby confirming that the corresponding ribosomal complex was a 70S IC carrying an fMet-tRNA^fMet^ at the P site and (ii) *E*_FRET_ vs. time trajectories had to exhibit at least one 0.55→0.31 transition after delivery of TCs, thereby confirming that the corresponding 70S IC had accommodated a *Pro-SufB2* or Pro-*ProL* into the A site, that the A site-bound Pro-*SufB2* or Pro-*ProL* had participated as the acceptor in peptide-bond formation, and that the resulting PRE complex was capable of undergoing GS1→GS2 transitions. We note that the second criterion might result in the exclusion of *E*_FRET_ vs. time trajectories in which Cy3 or Cy5 simply photobleached prior to undergoing a 0.55→0.31 transition, and could therefore result in a slight overestimation of *k*_70S IC→GS2_ and/or *k*_GS1→GS2_ (see below for a detailed description of how *k*_70S IC→GS2_, *k*_GS1→GS2_, and other kinetic and thermodynamic parameters were estimated). Nonetheless, the number of such *E*_FRET_ vs. time trajectories should be exceedingly small. This is because the rates with which the fluorophore that photobleached the fastest, Cy5, entered into the photobleached state (Ø) from the GS1, GS2, EF-G-bound GS2-like, and POST states were *k*_GS1→ø_ = 0.04 ± 0.02 s^-1^, *k*_GS2→ø_ = 0.07 ± 0.01 s^-1^, *k*_ss2(G)→ø_ = 0.07 ± 0.01 s^-1^ (where the subscript “(G)” denotes experiments performed in the presence of EF-G), and *k*_posτ→ø_ 0.05 ± 0.02 s^-1^, respectively (see below for a detailed description of how *k*_GS1→ø_, *k*_GS2→ø_, *k*_ss2(G)→ø_, and *k*_posτ→ø_ were estimated). These rates are, on average, about 11-fold lower than those of *k*_70s IC→GS2_ and *k*_GS1 →GS2_ (0.3–0.6 s^-1^ and 0.58–0.82 s^-1^ (Supplementary Table 4)). Consequently, we do not expect the measurements of *k*_70S IC→GS2_ and *k*_GS1→GS2_ to be limited by Cy3 or Cy5 photobleaching. Additionally, even if *k*_70S IC→GS2_ and *k*_GS1→GS2_ were slightly overestimated, they would be expected to be equally overestimated for *SufB2*- and *ProL* ribosomal complexes given that the rate of photobleaching would be expected to be very similar for *SufB2-* and *ProL* ribosomal complexes. Furthermore, because we are primarily concerned with the relative values of *k*_70S IC→GS2_ and *k*_GS1→GS2_ for *SufB2-* vs. *ProL* ribosomal complexes, rather than with the absolute values of *k*_70S IC→GS2_ and *k*_GS1→GS2_ for the *SufB2-* and *ProL* ribosomal complexes, such slight overestimations do not affect the conclusions of the work presented here.

To calculate *k*_70S IC→GS2_ and the corresponding error from the pre-steady-state experiments, we analyzed the 70S IC survival probabilities (Supplementary Figure 4, Tables 4 and 5)^37,67^. Briefly, for each trajectory, we extracted the time interval during which we were waiting for the 70S IC to undergo a transition to GS2 and used these ‘waiting times’ to construct a 70S IC survival probability distribution, as shown in Supplementary Figure 4. All 70S IC survival probability distributions were best described by a single exponential decay function of the type

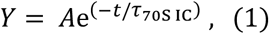

where *Y* is survival probability, *A* is the initial population of 70S IC, *t* is time, and *τ_70S IC_* is the time constant with which 70S IC transitions to a PRE complex in the GS2 state. *k*_70S IC→GS2_ was then calculated using the equation *k*_70S IC→GS2 = 1 / τ70S IC_. Errors were calculated as the standard deviation of technical triplicates.

Six sets of kinetic and/or thermodynamic parameters were calculated from hidden Markov model (HMM) analyses of the recorded movies. These parameters are defined here as: (i) *k*_GS1→GS2_, *k*_GS2→GS1_, and *K*_eq_ from the pre-steady-state and steady-state movies recorded for the delivery of *SufB2*- and *ProL*-TCs in the absence of EF-G (Figures 6d, 6f, and Supplementary Figure 3 and Table 4); (ii) *k*_GS2→POST_ from the pre-steady-state movie recorded for the delivery of *ProL*-TC in the presence of EF-G (Figures 6e, 6g, and Supplementary Figure 5 and Table 5); (iii) the fractional population of the POST complex from the pre-steady-state and steady-state movies recorded for the delivery of *SufB2*- and *ProL*-TCs in the presence of EF-G (Figures 6e, 6g, and Supplementary Figure 5 and Table 5); (iv) *K*_GS1→GS2_, *K*_GS2→GS1_, and *K*_eq_ from a sub-population of PRE complexes that lacked an A site-bound, deacylated *SufB2* in the steady-state movies recorded for the longer time points (i.e., 3, 10, and 20 min) after the delivery of *SufB2*-TC in the presence of EF-G (Figures 6g, Supplementary Table 6); (v) *K*_GS1→GS2_, *K*_GS2→GS1_, and *K*_eq_ from the steady-state movies recorded for the *SufB2-* and *ProL* PRE^-A^ complexes (Figures 6h and Supplementary Figure 6 and Table 7); and (vi) *K*_GS1→ø_, *K*_GS2→ø_, *k*_GS2(G)→ø_, and *k*_posτ→ø_ from the movies described in (i)-(v) (Figures 6d–6h, Supplementary Figures 3, 5, and 6, and reported two paragraphs above). To calculate these parameters, we extended the variational Bayes approach we introduced in the vbFRET algorithm^68^ to estimate a ‘consensus’ (*i.e.*, ‘global’) HMM of the *E*_FRET_ vs. time trajectories. In this approach, we use Bayesian inference to estimate a single, consensus HMM that is most consistent with all the *E*_FRET_ vs. time trajectories in a movie, rather than to estimate a separate HMM for each trajectory in the movie. To estimate such a consensus HMM, we assume each trajectory is independent and identically distributed, thereby enabling us to perform the inference using the likelihood function

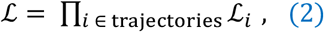

where 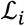 is the variational approximation of the likelihood function for a single trajectory. Subsequently, the single, consensus HMM that is most consistent with all of the trajectories is estimated using the expectation-maximization algorithm that we have previously described^68^. Viterbi paths (Supplementary Figures 3, 5, and 6), representing the most probable hidden-state trajectory, were then calculated from the HMM using the Viterbi algorithm^69^. Based on extensive smFRET studies of translation elongation using the bL9(Cy3)-uL1(Cy5) smFRET signal^35,36,38^, we selected a consensus HMM composed of three states for further analysis of the data. For calculation of the kinetic and/or thermodynamic parameters in (i), (iv), and (v), the three states corresponded to GS1, GS2, and ∅ and for calculation of the kinetic and/or thermodynamic parameters in (ii) and (iii), the three states corresponded to EF-G-bound GS2-like, POST, and ∅. The transition matrix of the consensus HMM was then used to calculate *k*_GS1→GS2_ and *k*_GS2→GS1_ in (i), (iv), and (v); *k*_GS2→POST_ in (ii); *k*_GS1→ø_, *k*_GS2→ø_, *k*_GS2(G)→ø_, and *k*_posτ→ø_ in (vi); and the errors corresponding to each of these parameters. This transition matrix consists of a 3 x 3 matrix in which the off-diagonal elements correspond to the number of times a transition takes place between each pair of the GS1, GS2, and ∅ states (in (i), (iv), (v), and (vi)) or each pair of the EF-G-bound GS2-like, POST, and ∅ states (in (ii) and (vi)) and the on-diagonal elements correspond to the number of times a transition does not take place out of the GS1, GS2, and ∅ states (in (i), (iv), (v), and (vi)) or out of the EF-G-bound GS2-like, POST, and ∅ states (in (ii) and (vi)). Each element of this matrix parameterizes a Dirichlet distribution, from which we calculated the mean and the square root of the variance for four transition probabilities *p*_GS1→GS2_, *p*_GS2→GS1_, *P*_GS1→ø_, and *P*_GS2→ø_ (in (i), (iv), (v), and (vi)) or for three transition probabilities *p*_GS2→POST_, *P*_GS2(G)→ø_, and *p*_posτ→ 0_ (in (ii) and (vi)). These transition probabilities were then used to calculate the corresponding four rate constants, *k*_GS1→GS2_, *k*_GS2→GS1_, *k*_GS1→ø_, and *k*_GS2→ø_ (in (i), (iv), (v), and (vi)) or three rate constants, *k*_GS2→POST_, *k*_GS2(G)→ø_, and *k*_posτ→ø_ (in (ii) and (vi)) using the equation

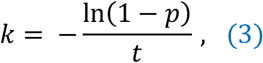

where *t* is the time interval between data points (*t* = 0.1 s). We propagated the error for the transition probabilities into the error for the rate constants using the equation

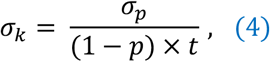

where *σ_p_* is the standard deviation of the variance of *p* and *σ_k_* is the standard deviation of the variance of *k*. *K*_eq_ in (i), (iv), and (v) was determined using the equation *K*_eq_ = *K*_GS1→GS2_ / *k*_GS2→GS1_. The fractional populations of the POST complex in (iii) and the corresponding errors were calculated by marginalizing, which in this case simply amounts to calculating the mean and the standard error of the mean, for the conditional probabilities of each *E*_FRET_ data point given each hidden state. Because the data points preceding the initial 70S IC→GS2 transition in the presteady-state movies do not contribute to the kinetic and/or thermodynamic parameters in (i)-(vi), these data points were not included in the analyses that were used to determine these thermodynamic parameters.

## QUANTIFICATION AND STATISTICAL ANALYSES

All ensemble biochemical experiments and cell-based reporter assays were repeated at least three times and the mean values and standard deviations for each experiment or assay are reported. Technical replicates of all smFRET experiments were repeated at least three times and trajectories from all of the technical replicates for each experiment were combined prior to generating the surface contour plot of the time evolution of population FRET and modeling with the HMM. Mean values and errors for the transition rates and fractional populations determined from modeling with an HMM are reported (for details see “Analysis of smFRET experiments” in Methods). Mean values and standard deviations for the *k*_70S IC→GS2S_ were determined from technical triplicates of the survival plots analysis for each experiment and are reported.

## DATA AND CODE AVAILABILITY

### Data Availability

With the exception of the smFRET data, all other data supporting the findings of this study are presented within this article. Due to the lack of a public repository for smFRET data, the smFRET data supporting the findings of this study are available from the corresponding authors upon request. Source data are provided with this paper.

## Code Availability

The code used to analyze the TIRF movies in this study is described in a manuscript in preparation (Jason Hon, Colin Kinz-Thompson, Ruben L. Gonzalez), where R.L.G. is the corresponding author. Therefore, the code is available from R.L.G, upon request.

## ACKNOWLEDGEMENTS

We thank Dr. Hajime Tokuda for rabbit polyclonal anti-LolB antibodies, Dr. Colin Kinz-Thompson and Korak Kumar Ray for help with smFRET data analysis. R.L.G. and H.L. thank the Columbia University Precision Biomolecular Characterization Facility for access to and support of instrumentation. This work was supported by NIH grants GM134931 to Y-M.H. and GM119386 to R.L.G., a Charles H. Revson Foundation Postdoctoral Fellowship in Biomedical Science 19-24 to H.L., a Japanese JSPS overseas postdoctoral fellowship to I.M., and NSF grant CHE-1708759 to E.J.P.

## AUTHOR CONTRIBUTIONS

H.G. conceived of and performed ensemble rapid kinetic assays, R.L.G. and H.L. conceived of and designed smFRET assays, H.L. performed smFRET assays, I.M. performed cell-based reporter assays, D.M.R. and E.J.P. generated aminoacyl-DBE derivatives, T.C. performed G37 methylation and aminoacylation assays, and A.B.C. and G.B. provided *E. coli* 70S ribosomes. Y.M.H. and R.L.G. wrote the manuscript.

## COMPETING FINANICAL INTERESTS

The authors declare no competing interests.

## CONTACT FOR REAGENT AND RESOURCE SHARING

Further information and requests for resources and reagents should be directed to and will be fulfilled by the lead contacts Ruben L. Gonzalez, Jr. and Ya-Ming Hou.

## REFERENCES

1. Wang, K., Schmied, W.H. & Chin, J.W. Reprogramming the genetic code: from triplet to quadruplet codes. Angew Chem Int Ed Engl 51, 2288–97 (2012).

2. Chen, Y. et al. Controlling the Replication of a Genomically Recoded HIV-1 with a Functional Quadruplet Codon in Mammalian Cells. ACS Synth Biol 7, 1612–1617 (2018).

3. Lee, B.S., Kim, S., Ko, B.J. & Yoo, T.H. An efficient system for incorporation of unnatural amino acids in response to the four-base codon AGGA in Escherichia coli. Biochim Biophys Acta 1861, 3016–3023 (2017).

4. Chatterjee, A., Lajoie, M.J., Xiao, H., Church, G.M. & Schultz, P.G. A bacterial strain with a unique quadruplet codon specifying non-native amino acids. Chembiochem 15, 1782–6 (2014).

5. Niu, W., Schultz, P.G. & Guo, J. An expanded genetic code in mammalian cells with a functional quadruplet codon. ACS Chem Biol 8, 1640–5 (2013).

6. Wang, N., Shang, X., Cerny, R., Niu, W. & Guo, J. Systematic Evolution and Study of UAGN Decoding tRNAs in a Genomically Recoded Bacteria. Sci Rep 6, 21898 (2016).

7. Neumann, H., Wang, K., Davis, L., Garcia-Alai, M. & Chin, J.W. Encoding multiple unnatural amino acids via evolution of a quadruplet-decoding ribosome. Nature 464, 441–4 (2010).

8. Wang, K. et al. Optimized orthogonal translation of unnatural amino acids enables spontaneous protein double-labelling and FRET. Nat Chem 6, 393–403 (2014).

9. Atkins, J.F., Loughran, G., Bhatt, P.R., Firth, A.E. & Baranov, P.V. Ribosomal frameshifting and transcriptional slippage: From genetic steganography and cryptography to adventitious use. Nucleic Acids Res 44, 7007–78 (2016).

10. Atkins, J.F. & Bjork, G.R. A gripping tale of ribosomal frameshifting: extragenic suppressors of frameshift mutations spotlight P-site realignment. Microbiol Mol Biol Rev 73, 178–210 (2009).

11. Roth, J.R. Frameshift suppression. Cell 24, 601–2 (1981).

12. Bossi, L. & Roth, J.R. Four-base codons ACCA, ACCU and ACCC are recognized by frameshift suppressor sufJ. Cell 25, 489–96 (1981).

13. Qian, Q. et al. A new model for phenotypic suppression of frameshift mutations by mutant tRNAs. Mol Cell 1, 471–82 (1998).

14. Weiss, R.B., Dunn, D.M., Shuh, M., Atkins, J.F. & Gesteland, R.F. E. coli ribosomes re-phase on retroviral frameshift signals at rates ranging from 2 to 50 percent. New Biol 1, 159–69 (1989).

15. Jager, G., Nilsson, K. & Bjork, G.R. The phenotype of many independently isolated +1 frameshift suppressor mutants supports a pivotal role of the P-site in reading frame maintenance. PLoS One 8, e60246 (2013).

16. Fagan, C.E., Maehigashi, T., Dunkle, J.A., Miles, S.J. & Dunham, C.M. Structural insights into translational recoding by frameshift suppressor tRNASufJ. RNA 20, 1944–54 (2014).

17. Maehigashi, T., Dunkle, J.A., Miles, S.J. & Dunham, C.M. Structural insights into +1 frameshifting promoted by expanded or modification-deficient anticodon stem loops. Proc Natl Acad Sci U S A 111, 12740–5 (2014).

18. Dunham, C.M. et al. Structures of tRNAs with an expanded anticodon loop in the decoding center of the 30S ribosomal subunit. RNA 13, 817–23 (2007).

19. Hong, S. et al. Mechanism of tRNA-mediated +1 ribosomal frameshifting. Proc Natl Acad Sci U S A 115, 11226–11231 (2018).

20. Sroga, G.E., Nemoto, F., Kuchino, Y. & Bjork, G.R. Insertion (sufB) in the anticodon loop or base substitution (sufC) in the anticodon stem of tRNA(Pro)2 from Salmonella typhimurium induces suppression of frameshift mutations. Nucleic Acids Res 20, 3463–9 (1992).

21. Caliskan, N., Katunin, V.I., Belardinelli, R., Peske, F. & Rodnina, M.V. Programmed −1 frameshifting by kinetic partitioning during impeded translocation. Cell 157, 1619–31 (2014).

22. Taylor, D.J. et al. Structures of modified eEF2 80S ribosome complexes reveal the role of GTP hydrolysis in translocation. EMBO J 26, 2421–31 (2007).

23. Khade, P.K. & Joseph, S. Messenger RNA interactions in the decoding center control the rate of translocation. Nat Struct Mol Biol 18, 1300–2 (2011).

24. Liu, G. et al. EF-G catalyzes tRNA translocation by disrupting interactions between decoding center and codon-anticodon duplex. Nat Struct Mol Biol 21, 817–24 (2014).

25. Abeyrathne, P.D., Koh, C.S., Grant, T., Grigorieff, N. & Korostelev, A.A. Ensemble cryo-EM uncovers inchworm-like translocation of a viral IRES through the ribosome. Elife 5, doi: 10.7554/eLife.14874 (2016).

26. Schuwirth, B.S. et al. Structures of the bacterial ribosome at 3.5 A resolution. Science 310, 827–34 (2005).

27. Pulk, A. & Cate, J.H. Control of ribosomal subunit rotation by elongation factor G. Science 340, 1235970 (2013).

28. Ratje, A.H. et al. Head swivel on the ribosome facilitates translocation by means of intra-subunit tRNA hybrid sites. Nature 468, 713–6 (2010).

29. Gamper, H.B., Masuda, I., Frenkel-Morgenstern, M. & Hou, Y.M. Maintenance of protein synthesis reading frame by EF-P and m(1)G37-tRNA. Nat Commun 6, 7226 (2015).

30. Masuda, I. et al. tRNA Methylation Is a Global Determinant of Bacterial Multi-drug Resistance. Cell Syst 8, 302–314 e8 (2019).

31. Christian, T. & Hou, Y.M. Distinct determinants of tRNA recognition by the TrmD and Trm5 methyl transferases. J Mol Biol 373, 623–32 (2007).

32. Murakami, H., Ohta, A., Ashigai, H. & Suga, H. A highly flexible tRNA acylation method for non-natural polypeptide synthesis. Nat Methods 3, 357–9 (2006).

33. Walker, S.E. & Fredrick, K. Recognition and positioning of mRNA in the ribosome by tRNAs with expanded anticodons. J Mol Biol 360, 599–609 (2006).

34. Gamper, H.B., Masuda, I., Frenkel-Morgenstern, M. & Hou, Y.M. The UGG Isoacceptor of tRNAPro Is Naturally Prone to Frameshifts. Int J Mol Sci 16, 14866–83 (2015).

35. Fei, J. et al. Allosteric collaboration between elongation factor G and the ribosomal L1 stalk directs tRNA movements during translation. Proc Natl Acad Sci U S A 106, 15702–7 (2009).

36. Ning, W., Fei, J. & Gonzalez, R.L., Jr. The ribosome uses cooperative conformational changes to maximize and regulate the efficiency of translation. Proc Natl Acad Sci U S A 111, 12073–8 (2014).

37. Fei, J., Kosuri, P., MacDougall, D.D. & Gonzalez, R.L., Jr. Coupling of ribosomal L1 stalk and tRNA dynamics during translation elongation. Mol Cell 30, 348–59 (2008).

38. Fei, J., Richard, A.C., Bronson, J.E. & Gonzalez, R.L., Jr. Transfer RNA-mediated regulation of ribosome dynamics during protein synthesis. Nat Struct Mol Biol 18, 1043–51 (2011).

39. Boel, G. et al. The ABC-F protein EttA gates ribosome entry into the translation elongation cycle. Nat Struct Mol Biol 21, 143–51 (2014).

40. Chen, B. et al. EttA regulates translation by binding the ribosomal E site and restricting ribosome-tRNA dynamics. Nat Struct Mol Biol 21, 152–9 (2014).

41. Kim, H.K. et al. A frameshifting stimulatory stem loop destabilizes the hybrid state and impedes ribosomal translocation. Proc Natl Acad Sci U S A 111, 5538–43 (2014).

42. Munro, J.B., Wasserman, M.R., Altman, R.B., Wang, L. & Blanchard, S.C. Correlated conformational events in EF-G and the ribosome regulate translocation. Nat Struct Mol Biol 17, 1470–7 (2010).

43. Blanchard, S.C., Kim, H.D., Gonzalez, R.L., Jr., Puglisi, J.D. & Chu, S. tRNA dynamics on the ribosome during translation. Proc Natl Acad Sci U S A 101, 12893–8 (2004).

44. Studer, S.M., Feinberg, J.S. & Joseph, S. Rapid kinetic analysis of EF-G-dependent mRNA translocation in the ribosome. J Mol Biol 327, 369–81 (2003).

45. Wintermeyer, W. & Rodnina, M.V. Translational elongation factor G: a GTP-driven motor of the ribosome. Essays Biochem 35, 117–29 (2000).

46. Ermolenko, D.N. et al. Observation of intersubunit movement of the ribosome in solution using FRET. J Mol Biol 370, 530–40 (2007).

47. Ermolenko, D.N. & Noller, H.F. mRNA translocation occurs during the second step of ribosomal intersubunit rotation. Nat Struct Mol Biol 18, 457–62 (2011).

48. Cornish, P.V. et al. Following movement of the L1 stalk between three functional states in single ribosomes. Proc Natl Acad Sci U S A 106, 2571–6 (2009).

49. Nguyen, H.A., Hoffer, E.D. & Dunham, C.M. Importance of a tRNA anticodon loop modification and a conserved, noncanonical anticodon stem pairing in tRNACGGProfor decoding. J Biol Chem 294, 5281–5291 (2019).

50. Guo, Z. & Noller, H.F. Rotation of the head of the 30S ribosomal subunit during mRNA translocation. Proc Natl Acad Sci U S A 109, 20391–4 (2012).

51. Zhou, J., Lancaster, L., Donohue, J.P. & Noller, H.F. Spontaneous ribosomal translocation of mRNA and tRNAs into a chimeric hybrid state. Proc Natl Acad Sci U S A 116, 7813–7818 (2019).

52. Korniy, N., Samatova, E., Anokhina, M.M., Peske, F. & Rodnina, M.V. Mechanisms and biomedical implications of −1 programmed ribosome frameshifting on viral and bacterial mRNAs. FEBS Lett 593, 1468–1482 (2019).

53. Lajoie, M.J. et al. Genomically recoded organisms expand biological functions. Science 342, 357–60 (2013).

54. Wang, K., de la Torre, D., Robertson, W.E. & Chin, J.W. Programmed chromosome fission and fusion enable precise large-scale genome rearrangement and assembly. Science 365, 922–926 (2019).

55. Mohan, S., Donohue, J.P. & Noller, H.F. Molecular mechanics of 30S subunit head rotation. Proc Natl Acad Sci U S A 111, 13325–30 (2014).

56. Kaledhonkar, S. et al. Late steps in bacterial translation initiation visualized using time-resolved cryo-EM. Nature 570, 400–404 (2019).

57. Chen, B. et al. Structural dynamics of ribosome subunit association studied by mixing-spraying time-resolved cryogenic electron microscopy. Structure 23, 1097–105 (2015).

58. Reinkemeier, C.D., Girona, G.E. & Lemke, E.A. Designer membraneless organelles enable codon reassignment of selected mRNAs in eukaryotes. Science 363(2019).

59. Datsenko, K.A. & Wanner, B.L. One-step inactivation of chromosomal genes in Escherichia coli K-12 using PCR products. Proc Natl Acad Sci U S A 97, 6640–5 (2000).

60. Fei, J. et al. A highly purified, fluorescently labeled in vitro translation system for single-molecule studies of protein synthesis. Methods Enzymol 472, 221–59 (2010).

61. Christian, T., Lahoud, G., Liu, C. & Hou, Y.M. Control of catalytic cycle by a pair of analogous tRNA modification enzymes. J Mol Biol 400, 204–17 (2010).

62. Zhang, C.M., Perona, J.J., Ryu, K., Francklyn, C. & Hou, Y.M. Distinct kinetic mechanisms of the two classes of Aminoacyl-tRNA synthetases. J Mol Biol 361, 300–11 (2006).

63. Peacock, J.R. et al. Amino acid-dependent stability of the acyl linkage in aminoacyl-tRNA. RNA 20, 758–64 (2014).

64. Aitken, C.E., Marshall, R.A. & Puglisi, J.D. An oxygen scavenging system for improvement of dye stability in single-molecule fluorescence experiments. Biophys J 94, 1826–35 (2008).

65. Gonzalez, R.L., Jr., Chu, S. & Puglisi, J.D. Thiostrepton inhibition of tRNA delivery to the ribosome. RNA 13, 2091–7 (2007).

66. Desai, B.J. & Gonzalez, R.L., Jr. Multiplexed, bioorthogonal labeling of multicomponent, biomolecular complexes using genomically encoded, non-canonical amino acids. bioRxiv doi: 10.1101/730465(2019).

67. MacDougall, D.D. & Gonzalez, R.L., Jr. Translation initiation factor 3 regulates switching between different modes of ribosomal subunit joining. J Mol Biol 427, 1801–18 (2015).

68. Bronson, J.E., Fei, J., Hofman, J.M., Gonzalez, R.L., Jr. & Wiggins, C.H. Learning rates and states from biophysical time series: a Bayesian approach to model selection and single-molecule FRET data. Biophys J 97, 3196–205 (2009).

69. Viterbi, A.J. Error bounds for convolutional codes and an asymptotically optimum decoding algorithm. IEEE Trans. Inform. Theory 13, 260–269 (1967).

70. Agirrezabala, X. et al. Visualization of the hybrid state of tRNA binding promoted by spontaneous ratcheting of the ribosome. Mol Cell 32, 190–7 (2008).

